# The neural circuitry of emotion-induced distortions of trust

**DOI:** 10.1101/129130

**Authors:** Jan B. Engelmann, Friederike Meyer, Christian C. Ruff, Ernst Fehr

## Abstract

Aversive emotions are likely to be a key source of irrational human decision-making but still little is known about the neural circuitry underlying emotion-cognition interactions during social behavior. Here, we show that incidental aversive emotions distort trust decisions and cause significant changes in the associated neural circuitry. Experimentally-induced negative affect reduced trust, suppressed trust-specific activity in left temporoparietal junction (TPJ), and reduced functional connectivity between TPJ and emotion-related regions such as the amygdala. The posterior superior temporal sulcus (pSTS) seems to play a key role in mediating the impact of emotion on behavior: Functional connectivity of this brain area with left TPJ was associated with trust in the absence of negative emotions, but aversive emotions disrupted this association between TPJ-pSTS connectivity and behavioral trust. Our findings may be useful for a better understanding of the neural circuitry of affective distortions and may help identify the neural bases of psychiatric diseases that are associated with emotion-related psychological and behavioral dysfunctions.

---

Trust pervades almost every aspect of human social life. It plays a decisive role in families, organizations, markets and in the political sphere. Without trust, families fall apart, organizations are inefficient, market transactions are costly and political leaders lack public support. Research in behavioral economics and neuroeconomics has begun to elucidate the determinants and neural correlates of trust ^1-3^. However, despite recent progress in understanding the determinants of trust ^4^ and its distortions in psychiatric disorders ^5^, there are still large gaps in our knowledge about the impact of our emotions on trust ^6,7^, and particularly the underlying neural circuitry.

We focus here on incidental emotions, which have been shown to distort choice behavior for financial and other types of social decisions ^8-11^. Incidental emotions are of particular interest due to their ubiquity in real life and because they are prime candidates for emotion-induced behavioral distortions. By definition, incidental emotions are unrelated to choice outcomes and, to the extent to which they affect behavior, may cause irrational behavioral biases. Prominent theoretical accounts^12-14^ distinguish such incidental affect from anticipatory emotions that reflect how decision-makers expect to feel about the outcomes of their decisions. While recent research has made much progress in outlining the neural underpinnings of emotional processes on the one hand ^15,16^ and of decision-making on the other ^17,18^, the neural interactions between emotional and cognitive processes that support choice have largely been explored from theoretical perspectives^19-21^. It is therefore important to directly examine the behavioral and neural mechanisms by which incidental emotions distort decisions to trust.

To study the behavioral impact and the underlying neural circuitry of emotion-induced distortions of trust, we employed a modified version of the well-established trust game ^22^ that has also been used in a number of previous imaging studies^1,23-27^. In the trust game, two anonymous players, which we call investor and trustee, sequentially send money to each other. In the first stage, the investor faces the choice of whether and how much of her endowment to transfer to the trustee. Then the experimenter triples the sent amount, before it is transferred to the trustee. The investor’s decision to transfer money thus increases the total amount of money that can be distributed among the two players. In the second stage, the trustee is informed about the total amount that he received and then needs to decide how much of this money to send back (nothing, parts of it, or all of it). Thus, while the investor’s transfer increases the total amount of money available to both parties, the investor also faces the risk of not benefitting at all from the transfer because the trustee is completely free in his back-transfer decision. Therefore, the decision to transfer money constitutes an act of trust, as the investor makes herself vulnerable to the potentially selfish behavior of the trustee ^4^.

Trust decisions involve both a financial risk due to the possibility of losing the invested money, as well as a social risk of being betrayed by an untrustworthy trustee^2,3,28^. The latter social risk is indeed specific to trust, as compared to non-social types of risky choices. To enable clear identification of the impact of incidental emotion on the mechanisms specifically involved in trust, it is important to include a well-matched non-social control task that has equivalent financial payouts without the social risk of betrayal. For this reason, our subjects also faced a non-social control condition that was identical to the trust condition in every respect, except that instead of a trustee, a computer made a “back-transfer” that determined the profitability of the investor’s “transfer” ^25,29^. This back-transfer was determined using an algorithm that drew random samples from the probability distribution of the trustees’ decisions in the trust condition (see online Methods). Thus, the choice options and the profitability of the investor’s transfer in the non-social control and trust condition were exactly the same, and the distinguishing feature between the trust and the non-social control game was the unique possibility of betrayal by the interaction partner in the trust game ^2^. This difference between the trust and the non-social control game was saliently indicated at the beginning of each respective trial with either a human-like symbol on the computer screen (in the trust game) or a non-human symbol (in the non-social control game).

The unique possibility of being betrayed by the trustee provides a strong incentive to avoid such betrayal ^2^. Therefore, the investor needs to assess the likelihood of such betrayal when deciding how much money to entrust the interaction partner. This assessment of betrayal likelihood is accomplished by mentalizing, i.e., taking the perspective of the trustee to estimate how she will react to given transfers. In contrast, no such processes are necessary to make a decision in the non-social control task. Recent theoretical accounts ^30^ and neuroimaging studies ^23,31-34^ have confirmed that decisions to trust may indeed require mentalizing, since brain areas commonly found to be involved in mentalizing (including DMPFC, STS, TPJ) are activated during the trust game. We therefore expected that aversive emotions may exert their effect on decisions to trust by affecting trust-specific activation in regions involved in representing other people’s mental states, including temporoparietal junction (TPJ) and dorsomedial prefrontal cortex (DMPFC) ^5,35-37^.

To investigate the impact of aversive incidental emotion on trust decisions, subjects made decisions in either trust or non-social control trials within two different emotional contexts. They were either under the threat of relatively intense tactile stimulation that was somewhat painful (“threat condition”), or they faced the possibility of receiving weak tactile stimulation that was still noticeable, but clearly not painful. There was thus no anticipation of aversive events in this “no threat” condition (Figure 1a). A prolonged period of incidental aversive emotion was established by administering the tactile stimulations at unpredictable time points and frequencies for the duration of an entire block. A block consisted of several trust or control trials in the threat condition or several such trials in the no-threat condition. The threat-of-shock paradigm employed in the current study has been shown to reliably induce negative emotion ^38-40^ and addresses the limitations of standard emotion ^41,42^ and stress induction ^6,8,43,44^ procedures as follows: First, the threat of shock provides an immediate stimulus of biological significance that triggers an aversive and automatic emotional reaction, the intensity of which can be titrated to individual subjective percepts and measured throughout the experiment using standard psychophysiological techniques. Second, unlike standard emotion and stress inductions that are typically administered only once at the start of a session, threat-of-shock can be turned on and off throughout the duration of the entire experiment, thereby reinstating the emotional reaction at every presentation. This clearly disambiguates anxiety from stress recovery, which is hard to achieve with standard emotion and stress inductions ^45,46^. Third, threat of shock was administered within-subject in a single experimental session, therefore allowing each subject to serve as their own control. Finally, tactile stimulation was administered in both the trust and the non-social control condition, thus minimizing demand effects.

**Figure 1.**
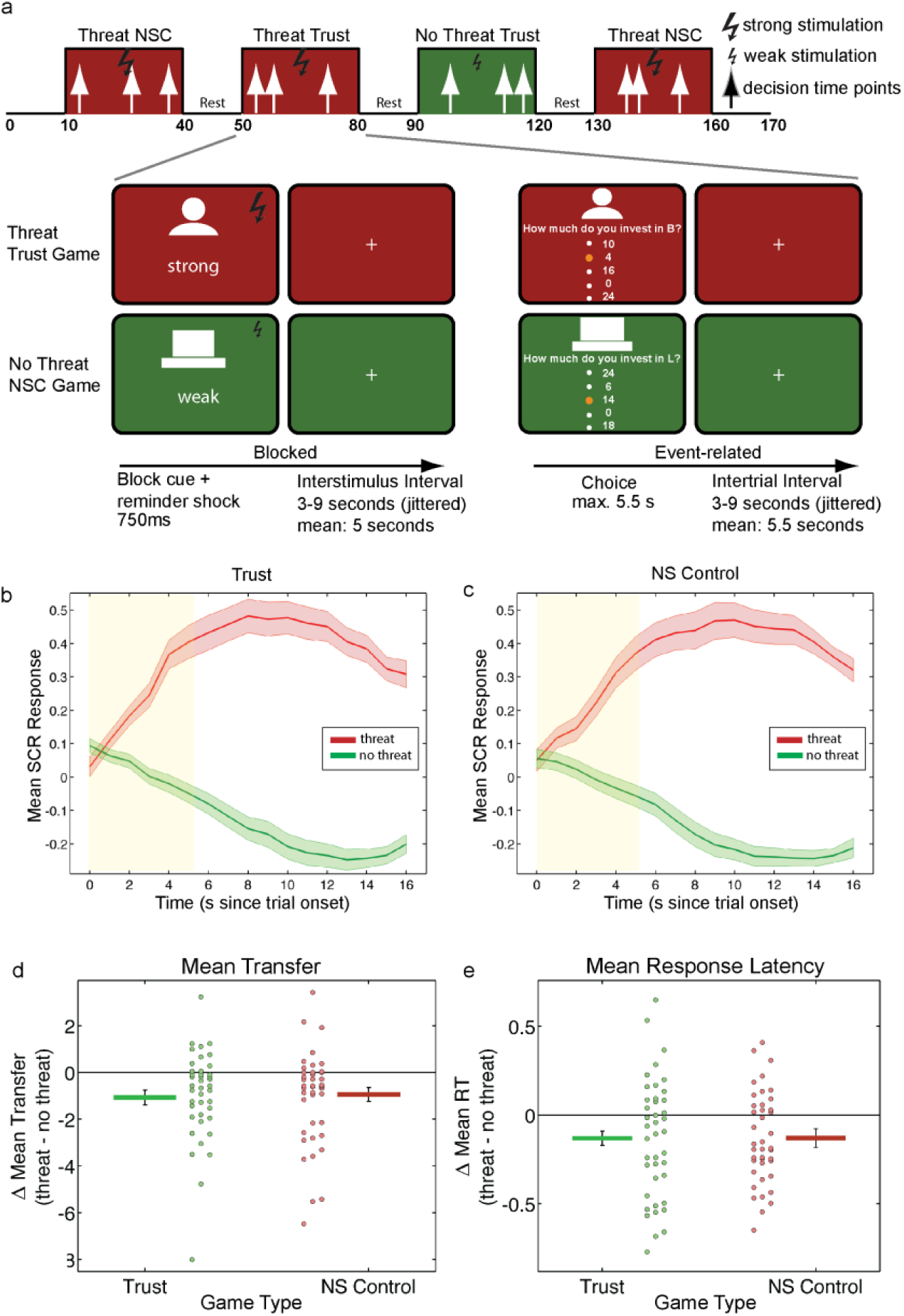
Experimental task, electrophysiological and behavioral findings. (a) Schematic representation of hybrid fMRI design, trial sequence and timing (see Methods). Subjects faced blocks of trust (human icon) and non-social control (NSC, computer icon) trials in random order. During trust and NSC blocks subjects expected either strong (“threat”) or weak (“no-threat”) tactile stimulation at unpredictable times. At the beginning of each block, a 750-ms visual cue followed by tactile stimulation reminded subjects of the game type (trust or non-social control) and stimulation intensity (weak or strong) for the current block. On each trial, subjects chose how much of their endowment of 24 CHF to transfer to a stranger (trust game), or invest in an ambiguous lottery that provided a 40-60% probability of returning an amount greater than the investment (NSC game). (b-c) The threat of an aversive tactile stimulation, not the shock itself (see Methods), leads to a strong increase in skin conductance responses (SCR) in (b) the trust game (p < 0.0001) and (c) the non-social control game (p < 0.0001), (d) In the threat condition (relative to the no-threat condition) subjects transferred significantly less to an anonymous stranger in the trust game (p < 0.005, reduction due to threat in 71% of subjects) and invested less into an ambiguous lottery in the non-social control game (p < 0.005, reduction due to threat in 73% of subjects). These results are driven by the emotional arousal induced by the threat of a shock and not by the actual experience of shocks shortly before choice (Table S1). In the threat condition (relative to the no-threat condition) subjects made their decisions significantly faster in both the trust (p < 0.005) and the control (p < 0.05) game. Dot plots reflect the difference between mean transfer (d) and response latency (e) in the threat compared to the no threat condition for the same subject to illustrate the *reduction* of mean transfer and response latency due to threat.

We conjectured that incidental aversive emotion modulates trust-specific computations, particularly the simulations of the trustee’s reaction to given transfers. This has potentially important implications because if incidental aversive emotion disrupts the recruitment of the social-cognitive processes necessary for mentalizing and perspective taking, we are likely to observe that aversive emotion also has specific effects on the neural mechanisms of trust decisions. We expected that incidental aversive emotion influences trust decisions by specifically modulating neural responses in regions involved in representing other people’s mental states, including temporoparietal junction (TPJ) and dorsomedial prefrontal cortex (DMPFC) ^5,35,47,48^. Three pieces of evidence support our neuroanatomical hypothesis. First, theoretically, investors are highly motivated to estimate the likelihood of betrayal via simulations of the trustee’s reaction to given transfers as a fundamental aspect of their decisions to trust. Previous research has consistently implicated TPJ and DMPFC in such simulations of others’ mental states (e.g., ^5,35,47,48^, also Figure S1). Second, previous research has demonstrated the involvement of the same neural structures (TPJ, DMPFC) also during trust decisions and similar social interaction ^23,27,34^. Third, a conjunction of the neurosynth meta-analyses for the terms “emotion” and “theory of mind” identifies an overlap between these networks in TPJ and DMPFC (Figure S1), therefore implicating these regions in both emotional and social cognitive processes. Together, the above results establish these regions as prime candidates for investigations of the modulatory effects of incidental aversive emotion on mentalizing during trust decisions. In addition, we explored the effects of incidental aversive emotion on neural activity in the anterior insula, which has consistently been implicated in both aversive emotion ^49,50^ and trust decisions ^51^, and the amygdala, which has consistently ^52^ and relatively specifically (neurosynth reverse inference analysis for “aversive”) been implicated in the processing ^15^ and regulation of aversive emotions ^53^, and at the same time plays a central role in trustworthiness inferences ^54,55^.

## Results

### Threat of shock induces autonomic arousal and aversive emotions during decision-making

We scanned 41 volunteers while they made trust decisions during the emotionally aversive threat condition and during the emotionally neutral no-threat condition. We first ascertained that threat of shock indeed induced emotional arousal by probing how galvanic skin conductance responses (SCR), self-reported emotion and brain activations changed in response to the threat of electrical stimulation (note that SCR was modeled with a general linear model that included regressors for the time points of actual shocks; the variance due to the shocks themselves was therefore removed from the analysis). As illustrated in Figures 1b and 1c, mean SCRs during both trust and non-social control trials were significantly greater during the threat condition compared to the no-threat condition [significant main effect of threat: F(1,39) = 141.27, p < 0.001, η^2^ = 0.784; significant main effect of time: F(16, 624) = 12.12, p < 0.001, η^2^ = 0.237; and, importantly, a significant two-way interaction between threat and time: F(16,624) = 99.28, p < 0.001, η^2^ = 0.718. The analysis yielded no other effects, all p > 0.5]. Follow-up pairwise comparisons at each time point showed significantly enhanced SCR during threat relative to no threat from 2 until 16 seconds after trial onset during trust, and from 3 until 16 seconds after trial onset during non-social control decisions (all two-tailed tests survive Bonferroni correction for multiple comparisons with all t(39) > 3.313 and all p < 0.002). Taken together, these results indicate significantly greater emotional arousal during the threat condition relative to the no-threat condition in both social and non-social game types.

The emotional arousal illustrated in Figures 1b and 1c was clearly experienced as aversive by the subjects. In an open-ended questionnaire administered after scanning, 95.12% of subjects responded that they experienced aversive emotional arousal during threat blocks (Supplementary Figure 1a). The aversive nature of the threat condition was further confirmed by strong activations of central nodes of the brain’s pain matrix during the (actual) experience of strong compared to weak tactile shocks (Supplementary Figure 1c) and by the observation of enhanced SCRs following the (actual) experience of strong compared to weak tactile shocks (Supplementary Figure 1b).

Jointly, the above electrophysiological results, self-reported emotions and activation within the brain’s pain matrix during and after the shock, indicate that subjects experienced the threat of a shock as an aversive and arousing emotional state. This state was clearly unrelated to the monetary outcome of trust- and risk-taking, as it did not affect the trustee’s or the computer’s decisions. The next question we addressed is whether this incidental emotional state distorts subjects’ behavior relative to the no-threat control condition.

### Aversive emotion reduces investments during trust decisions

To identify whether the aversive emotional state had a significant impact on decision-making, we first investigated mean transfer rates during trust and non-social control decisions for each emotional context and submitted these data to a two-way repeated-measures ANOVA (Mean transfer rates were normally distributed as indicated by the Shapiro-Wilk normality test: W = 0.976, p = 0.510) with the factors game type (trust, control) and threat (absent, present). Aversive emotional state significantly changed transfers during both trust and non-social control trials (Figure 1d), as indicated by a significant main effect of threat [F(1,40) = 17.483, p < 0.001, η^2^ = 0.304, see Supplementary Tables 2a and 2b for additional effects]. Moreover, separate pairwise comparisons (all two-tailed) showed that the threat condition led to a reduction of investments (Figure 1d) in the trust game [t(40) = -3.4, p < 0.005, mean transfer difference = -1.1 CHF] and in the non-social control game [t(40) = -3.16, p < 0.005, mean transfer difference = -0.93 CHF]. To exclude the possibility that choices were affected by the *actual experience* of shocks, rather than by the ongoing aversive emotion due to shock expectation, we ran several multiple regression analyses (Supplementary Text 1). The regression results (Supplementary Table 1) show that the behavioral results reported above were indeed due to the aversive emotion (p < 0.001) generated by the *threat* of shock, rather than reflecting the effect of actual shock experience immediately before decisions are taken (p = 0.23).

Aversive emotion also led to faster reaction times during both trust- and non-social control trials (Figure 1e). Mean reaction times were submitted to a two-way repeated-measures ANOVA (Mean reaction times were normally distributed as indicated by the Shapiro-Wilk normality test: W = 0.972, p = 0.402) with the factors game type (trust, control) and threat (absent, present). We obtained a significant main effect of threat [F(1,40) = 17.01, p < 0.001, η^2^ = 0.298, see Supplementary Tables 3a and 3b for additional effects]. The main effect of threat is characterized by significantly (two-tailed) faster mean reaction times in the threat relative to the no-threat condition (Figure 1e) for both the trust game [t(40) = -3.3, p < 0.005, mean RT difference = -0.13 s] and the non-social control task [t(40) = -2.5, p < 0.05, mean RT difference = -0.13 s].

Taken together, these behavioral results indicate that aversive emotion significantly reduced trust, as reflected by diminished transfer rates in the trust game. Additionally, aversive emotion reduced transfer rates in the nonsocial control task and reaction times in both the trust and the nonsocial control task. Notably, the absence of a significant interaction between threat and game type for electrophysiological and behavioral measures (transfer rates: F(1,40) = 0.122, p = 0.7, η^2^ = 0.003, response latencies: F(1,40) < 0.001, p = 0.993, η^2^ < 0.001, SCR: F(1,39) = 0.006, p = 0.938, η^2^ < 0.001) indicates that the impact of aversive emotion during trust and nonsocial control trials is similar across these multiple measurement modalities, confirming that our non-social condition constitutes a well-matched control for the trust game.

### Behavioral differences between trust and control decisions

Despite the similar effects of aversive emotion on transfer rates and reaction times across the two game types, three sources of evidence underline that subjects treated decisions in the trust and non-social control games differently and, therefore, that aversive emotion impacted separable underlying cognitive processes in the trust and non-social control game. First, average transfer rates were significantly higher in the trust (15.37 (0.767) MU) compared to the non-social control (12.671 (0.804) MU) game as indicated by the significant main effect of game type [F(1,40) = 20.319, p < 0.001, η^2^ = 0.337, see Supplementary Tables 2a and 2b for additional effects]. Moreover, average reaction times were significantly faster in the trust (2.575 (0.07) sec.) compared to the non-social control game (2.675 (0.066) sec.) as indicated by the significant main effect of game type [F(1,40) = 6.226, p = 0.017, η^2^ = 0.135, see Supplementary Tables 3a and 3b for additional effects]. Second, transfer rates within each game type were highly similar across threat conditions, but were significantly different across the two game types within each threat condition. That is, correlation analyses investigating relationships between the transfer rates within and across each game type showed stronger correlations between transfer rates for threat and no threat within each game type (Trust (threat, no threat): r = 0.931; NS control (threat, no threat): r = 0.937) and somewhat weaker correlations between transfer rates for trust and control decisions within each threat condition (threat (trust, risk): r = 0.739; No threat (trust, risk): r = 0.636). We compared correlations within (modified Pearson-Filon test, ZPF) and across (Williams t-test) game types using methods outlined in ^56^. This revealed that the correlations between transfer rates in the absence compared to the presence of threat within each game type were similar (ZPF = -0.23, p = 0.816). However, the correlation between transfer rates in the trust game during threat and no threat was significantly different from (1) the correlation between transfer rates under threat for trust and NS control games (t(38) = 4.23, p < 0.001) and from (2) the correlation between transfer rates under no threat for trust and NS control games (t(38) = 5.53, p < 0.001). Third, emotion-related changes in mean transfer rates for the trust game (no threat - threat) did not correlate significantly with emotion-related changes in transfer in the risk game (Pearson’s r = 0.231, p = 0.146). Together, these results indicate that subjects engaged in different cognitive processes to make decisions in the two game types, supporting the notion that the emotion-induced changes in the two game types reflect the influences of emotions on separable choice-related processes.

### Aversive emotion suppresses trust-related activity in TPJ

The main goal of our fMRI analyses was to identify the impact of aversive emotion on the neural mechanisms instantiating the social-cognitive processes that support trust decisions. We therefore first examined brain activation in the ROIs that we conjectured (see our hypotheses in the introductory section) to be preferentially engaged during *trust-specific* computations, such as the assessment of the trustee’s trustworthiness and the associated interplay between social cognition and social valuation. Regions involved in representing other people’s mental states include temporoparietal junction (TPJ) and dorsomedial prefrontal cortex (DMPFC) ^36^. We employed small-volume correction at an FWE-corrected threshold of p < 0.05 *in truly independent ROIs* defined with reverse inference maps from relevant search terms on neurosynth.org ^57^ (see online Methods). Where appropriate, we supplemented these hypothesis-driven ROI analyses with exploratory whole-brain analyses as outlined in the online methods. Furthermore, our fMRI analyses followed these steps: (1) we tested whether our a-priori regions of interest are involved in the processing of social-context or emotion by investigating main effects; (2) we then probed activity in these regions for the specific suppression of trust-related activity by threat by investigating interactions; (3) interaction results were then further characterized by simple comparisons within specific contexts (e.g., threat vs no-threat in the trust task). In the final part of this paper we also examined the domain general effects (i.e. the effects that are not specific to the trust game) of aversive emotions.

As a first step, our analyses confirmed that several of the conjectured regions were indeed specifically involved in trust (vs. non-social control) decisions (Main effect of task: Trust_no threat_ + Trust_threat_ > NS Control_no threat_ + NS Control_threat_). That is, an area in left TPJ (-57, -60, 27; k = 9; SV FWE-corrected, green region in Figure 2a) showed significantly greater activation during decision-making in trust relative to control trials in the absence of threat (activation SV FWE-corrected, see also Supplementary Table 2a). We then examined which of the ROIs showed a breakdown of trust-specific activity due to aversive emotion. To this end, we identified the threat-induced reduction in brain activation that was specific to the trust game with the following interaction contrast: (Trust_no threat_ > Trust_threat_) > (NS Control_no threat_ > NS Control_threat_). A region in left TPJ indeed showed this significant interaction effect (-60, -54, 19; k = 34, SV FWE-corrected, red region in Figure 2a, see also Supplementary Table 2b). To further characterize this interaction, we examined post hoc the impact of aversive emotion in the trust game separately from its impact in the non-social control game: This revealed a suppression of activation within left TPJ (-58, -55, 19; k = 35; SV FWE-corrected, Supplementary Table 2c) for trust decisions, but not during non-social control decisions (no voxels for NSC_No Threat_ > NSC_Threat_), even at a very liberal threshold of p < 0.05, uncorrected (see also Supplementary Figure 2a for additional univariate analyses that underline the strength of the interaction effect in left TPJ). These results indicate that the interaction effect is based on a selective interference of aversive emotion with trust-related activity, but not with activity in the nonsocial control condition. We also found a region in left anterior insula that showed the same interaction effect (-46, 14, -12, SV FWE-corrected, Supplementary Table 2b), but the only marginally significant simple effects (Trust_no threat_ > Trust_threat_: -46, 26, 12, k = 4, SV FWE-corrected p = 0.078; NSC_No Threat_ > NSC_Threat_: no voxels) indicate a weaker trust-related suppression of activity compared to TPJ.

**Figure 2.**
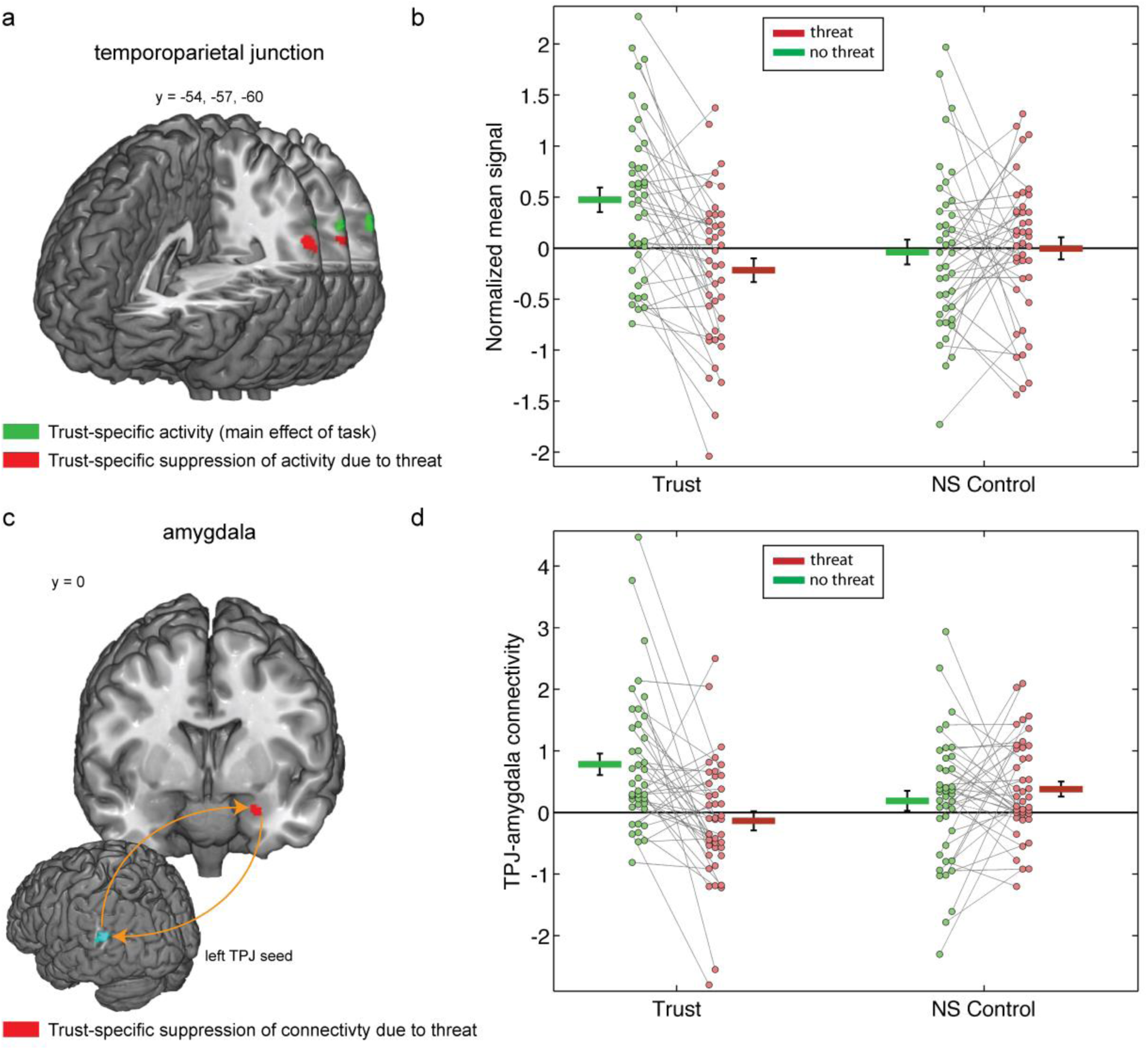
The impact of aversive emotion on trust-specific TPJ activity and connectivity. Panel (a) shows the region of left temporoparietal junction (peak at xyz = -57, -60, 27) that is selectively involved in trust compared to the non-social control task as reflected by a significant main effect of game type (shown in green, see also Table S2a). Importantly, aversive emotions induced by the threat of a shock reduced activation in left TPJ (relative to “no threat”) significantly more during the trust game than in the nonsocial control game (significant interaction effect, peak at xyz = -60, -54, 19, Table S2b). Voxels whose activity reflects this interaction effect are shown in red. All regions survive p<0.05 SVC-FWE-corrected (see methods). Threat-induced reduction of TPJ activity was observed in 78% of subjects during trust decisions (downward-sloping connecting lines), and in 44% of subjects during non-social control decisions, as shown in panel (b). The parameter estimates in (b) are extracted from a sphere (6-mm radius) around individual peaks within the TPJ cluster marked in red in panel a. (c) The left amygdala (peak at xyz = -28, -6, -14, see Table S3a) shows significantly stronger connectivity with TPJ during trust relative to control decisions as reflected by a significant main effect of threat. This coupling is disrupted by the threat of a shock specifically during trust as compared to the non-social control task (significant interaction effect; peak at xyz = -26, 0, -23, shown in red). All regions survive p<0.05 SVC-FWE-corrected (see methods). Threat-induced reduction of TPJ-amygdala connectivity was observed in 76% of subjects during trust decisions (downward-sloping connecting lines), and in 44% of subjects during non-social control decisions, as shown in panel (d). The parameter estimates are extracted from a 6-mm sphere around the individual peaks within the amygdala cluster marked in yellow in panel c, to visualize the specific effects of aversive emotion on functional connectivity between the left TPJ and left amygdala during decisions in the trust game. Dot plots in panels (e) and (d) reflect individual subject mean activation in each condition and are connected to illustrate the suppression of activity due to threat for each subject.

To identify to what extent the above effects are lateralized, we also inspected the main and interaction effects in right TPJ at uncorrected thresholds. This revealed that the right TPJ in fact also showed a numeric (but non-significant) main effect of trust (57, -55, 24, k = 45, p = 0.002) and a numeric, non-significant interaction effect in a superior (62, -54, 28, k = 11, p = 0.004) and an inferior (64, -45, 4, k = 15, p = 0.002) location in pSTS that is part of our TPJ mask. Thus, the effects we observed were somewhat more pronounced in the left hemisphere but not clearly lateralized.

Given that decisions involving trust rely on neural circuitry that mediates the interplay between social cognition and valuation ^1^, we also performed an exploratory analysis of the impact of aversive emotion on trust-related activity within regions commonly implicated in valuation (vmPFC and ventral striatum ^58^). No significant interactions were found, but tests of simple effects comparing threat and no threat during trust decisions showed reductions in trust-related activity due to aversive emotion in vmPFC and ventral striatum (Supplementary Table 2c).

### Aversive emotion suppresses trust-specific connectivity between TPJ and amygdala

Recent studies stress the importance of the interplay between cognitive and emotional networks ^15,59^. Therefore, we investigated the effects of aversive emotion on the connectivity between trust-relevant brain regions with Psychophysiological Interaction analyses (PPI) ^60^. In view of the key role of the temporoparietal junction (TPJ) in perspective taking and mentalizing ^5,47^ and the conjecture that these mental operations are important for trust, and our finding of enhanced activity in TPJ in the trust compared to the control task (see above), we were particularly interested in how aversive emotion changed the functional connectivity between the TPJ and emotion processing regions, such as the amygdala. ROI analysis of a Psychophysiological Interaction (PPI ^60^) analysis seeded in the TPJ confirmed that TPJ-amygdala connectivity was particularly disrupted by threat (No Threat_trust_ + No Threat_ns control_ > Threat_trust_ + Threat_ns control_) in left amygdala (-28, -6, -14; p < 0.05, SV FWE-corrected, k = 14, Supplementary Table 3a). Therefore, we were interested whether there was a *trust-specific* threat-induced connectivity change. To answer this question we performed an interaction analysis that examined whether the threat-induced connectivity change in the trust condition (Trust_no threat_ > Trust_threat_) is larger than the threat-induced connectivity change in the control condition (NS Control_no threat_ > NS Control_threat_).

This analysis revealed that threat-induced aversive emotion caused a connectivity change between TPJ and a region in the left amygdala [-26, 0, -23; p < 0.05, SV FWE-corrected, k = 12, Figure 2c shown in red] that was significantly larger in the trust game compared to the control task (see Supplementary Table 3b). We performed a post-hoc inspection of the significant interaction in left amygdala by investigating the effect of threat on connectivity changes for the trust and the non-social control condition separately. Aversive emotion disrupted functional connectivity specifically during trust (compare red vs green bars in Figure 2d) but not during non-social control decisions. A follow-up contrast investigating threat effects on trust-related connectivity patterns (Trust_No threat_ > Trust_Threat_) confirmed the suppression of TPJ-amygdala connectivity during trust decisions (-28, -6, -14; p < 0.05, SV FWE-corrected, k = 23, Supplementary Table 3c). In contrast, during decisions in the non-social control task, no voxels in our left amygdala ROI showed greater connectivity during no threat relative to threat, or the reverse contrast of threat vs. no threat, even at a very liberal threshold of p < 0.05, uncorrected (see also Supplementary Figure 2b for additional analyses that underline the strength of the interaction effect in left amygdala). Moreover, comparison of connectivity during decisions in the control compared to the trust task in the threat condition showed a significant suppression of connectivity during trust relative to non-social decisions in left amygdala (-26, 0, -23, p < 0.05, SV FWE-corrected, k = 14, Supplementary Table 3d), indicating that the threat-related suppression of TPJ-amygdala connectivity was also evident when comparing NS control to trust. Together, these results indicate that threat caused a specific suppression of connectivity during trust that can be observed when comparing the effect of threat within the trust task (Trust: no threat > threat), as well as the effect of threat on connectivity during trust relative to NS control decisions (Threat: ns control > trust). This suppression of TPJ-amygdala connectivity during trust decisions occurred in the absence of suppression during non-social control decisions (ns control: threat = no threat). Thus, aversive emotion not only affected trust-specific overall activation in the TPJ, but also led to trust-specific connectivity changes of this area with the amygdala.

To identify to what extent the functional connectivity effects between TPJ and amygdala were lateralized, we repeated the tests of main and interaction effects in right amygdala at uncorrected thresholds. We found a numeric but non-significant threat-induced disruption of TPJ-amygdala connectivity in the right amygdala (Main effect of threat: 21, -6, -12, k = 5, SV FWE-corrected p = 0.079). Furthermore, we found a numeric, non-significant interaction effect indicating trust-related disruption of TPJ-right amygdala connectivity (30, -7, -12, k = 2, p = 0.008). As for the TPJ, amygdala involvement was more pronounced for the left hemisphere but not clearly lateralized.

### A trust network: TPJ connectivity strength with pSTS, DMPFC and VLPFC specifically predicts behavioral trust

The above PPI analyses show the average impact of aversive emotion on the functional connectivity between TPJ and amygdala. However, as we observed strong individual differences in the functional connectivity between TPJ and amygdala on the one hand, and in mean transfer levels on the other hand, we asked the question how individual differences in functional TPJ connectivity are related to individuals’ mean transfer levels in the absence and the presence of aversive emotion. Following our analysis approach for TPJ activity above, we first identified trust-specific brain-behavior correlations by comparing the relationship between transfer rates and TPJ functional connectivity as a function of game type, regardless of the threat condition (Main effect of task: Trust_no threat_ + Trust_threat_ > NS Control_no threat_ + NS Control_threat_). Given the importance of social cognitive processes for trust decisions, we expected stronger TPJ connectivity with other regions implicated in social cognition, such as our a priori ROIs in DMPFC, TPJ and amygdala, for enhanced trust. We therefore examined in our ROIs whether the relationship between mean transfers and functional TPJ connectivity is different in the trust game compared to the non-social control game via a flexible factorial model that, in addition to the factors Subject, Task and Threat, also includes mean transfer levels in each condition as covariates. We indeed found a significantly stronger positive correlation with mean transfer rates in the trust game compared to the non-social control game for connectivity between the left TPJ and the right amygdala (28, 2, -20, k = 16), the DMPFC (-12, 54, 40, k = 30), the bilateral STS (right: 64, -43, 4, k = 30; left: -62, -52, -5, k = 10; note (see online methods) that the STS is the most ventral part of our TPJ mask), as well as bilateral anterior insula (left: -51, 21, -6, k = 245; right: 56, 18, 1, k = 525,) (all p < 0.05, SV FWE-corrected, Supplementary Table 4a).

For completeness, we also conducted an exploratory whole-brain analysis (FWE-corrected at the cluster level) that identified an extended network of regions (Figure 3a-b, d; Supplementary Table 5a) showing a difference in the relationship between individuals’ mean transfers and their functional TPJ connectivity across trust and control games. This analysis extends our ROI analysis by identifying regions outside our ROIs, including bilateral inferior frontal gyrus (right: 42, 27, -11, k = 1422, left: -51, 21, -6, k = 343) and bilateral dorsolateral PFC (right: 44, 8, 28, k = 601; left: -52, 6, 18, k = 415; see Supplementary Table 5a]. In all these regions we observed a positive and significantly stronger correlation between mean transfer levels and functional TPJ connectivity in the trust compared to the control task (see Figure 3a-b, d and Supplementary Table 5a).

**Figure 3.**
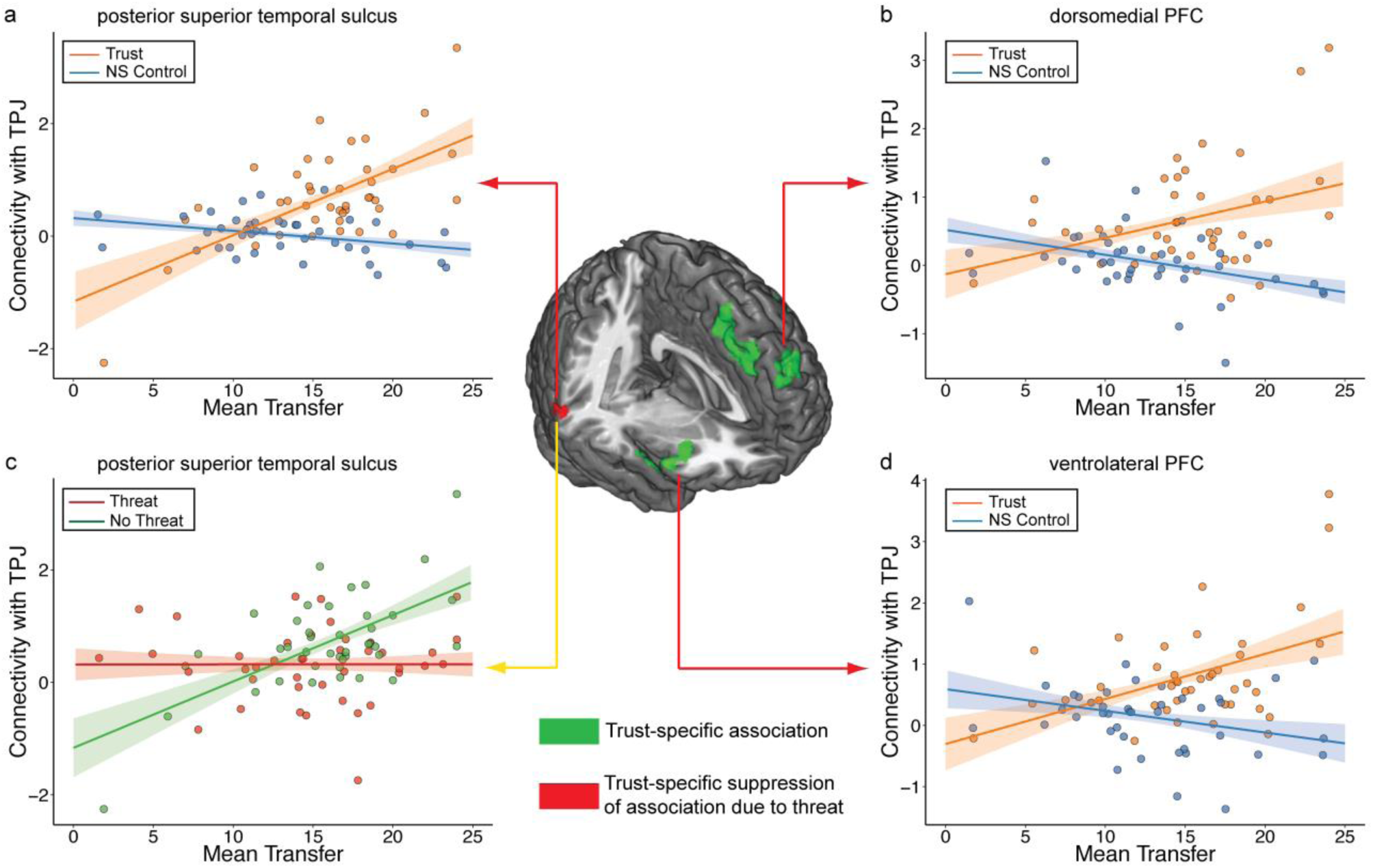
Trust-specific functional connectivity (a-b, d) and threat-induced breakdown of connectivity (c) between TPJ and a network of target regions. (a-b, d) Trust specific associations between transfer rate and trust-related neural activity reflecting the main effect of trust are shown in green activation clusters: Connectivity between left TPJ and its targets is positively associated with trust (orange regression lines) but not with non-social control decisions (blue regression lines) in (a) bilateral posterior superior temporal sulcus (left peak at xyz = -62, -52, -5; right peak at xyz = 64, -43, 4), (b) dorsomedial PFC (peak at xyz = -12, 54, 40), anterior insula (left peak at xyz = -51, 21, -6; right peak at xyz = 56, 18, 1) and amygdala (peak at xyz = 28, 2, -20). In contrast, mean transfers (i.e. investments) during the non-social (NS) control task (blue regression lines) are associated with reduced connectivity strength between TPJ and these regions. In all cases, the correlation between mean transfer and connectivity strength is stronger in the trust game compared to the non-social control task (see Table S4a for ROI analyses). (c) The results from the interaction contrast reflecting a trust-specific breakdown of the association between mean trust and TPJ connectivity is shown in the red activation cluster in STS (peak at xyz = 64, -43, 6, Table S4b): Aversive emotion causes a breakdown of the association between TPJ-pSTS connectivity and mean trust. The correlation between mean trust levels and TPJ-pSTS connectivity is stronger in the no threat compared to the threat condition (peak at xyz = 64, -43, 6; see Table S4c). Specifically, there is a positive association between TPJ-pSTS connectivity and the mean trust level when distortionary aversive emotion is absent (green regression line), which is eliminated by threat (red regression line). This suggests that connectivity between TPJ and its target region in pSTS supports general trust taking only in the absence of threat. The regression lines in (a-d) predict functional connectivity strength as a function of mean transfer levels based on an extended OLS model that estimates both the *slope* of the relationship between mean transfers and functional connectivity in the non-social control task and the *increase* in this relationship in the trust task (relative to the non-social control task). For this purpose we extracted the data from 6 mm spheres around individual interaction peak voxels (see online methods). Confidence bounds around regression lines reflect 95% confidence intervals around the model fit.

Finally, we also tested whether the slightly negative slopes observed in the regression lines connecting TPJ connectivity and mean transfer rates in the non-social control conditions of Figure 3a-b and 3d are statistically significant. To this end, we ran simple effects contrasts probing for a correlation between TPJ connectivity strength and mean transfer during NS control decisions in the absence of threat. We found no evidence that TPJ connectivity with its target regions negatively predicts transfer rates in the NS control condition, even at a relaxed threshold of p < 0.05. During trust decisions in the absence of threat, on the other hand, TPJ connectivity strength with pSTS (64, -43, 4, k = 335), dmPFC (20, 50, 39, k = 475), as well as left IFG (-51, 27, -3, k = 429), left posterior insula (-40, 3, 7, k = 578) and right IPS (44, -48, 40, k = 691) positively predicted transfer rates (Supplementary Table 5d).

Taken together, the above results therefore confirm the conjecture that aversive emotions suppress trust-specific functional connectivity between the TPJ and amygdala. In addition, the larger the TPJ connectivity with key regions implicated in mentalizing (the DMPFC and right STS) and emotion (the amygdala), the more subjects were willing to trust their partners on average. This predictive relationship between transfer rates and TPJ connectivity was absent in the non-social control game. These results thus suggest that trust involves functional communication between the TPJ and a network consisting of the amygdala, right STS, DMPFC and bilateral VLPFC.

### Aversive emotion removes the relationship between TPJ connectivity strength and behavioral trust

How did the relationship between functional connectivity patterns in the trust network and behavioral trust change if subjects were exposed to aversive emotion? Our previous results showing (1) greater TPJ connectivity with DMPFC and STS under enhanced trust (Fig. 3) but also, under conditions of threat (2) reduced behavioral trust (Fig. 1) and (3) suppressed trust-related TPJ activity (Fig. 2) jointly lead to the prediction that aversive emotion should have suppressive effects on the social cognitive machinery that supports trust decisions. We therefore expected a suppression of the relationship between behavioral trust and TPJ connectivity with other social cognition regions under conditions of threat. We tested this hypothesis in our a priori ROIs in TPJ and DMPFC. Specifically, we examined whether there was a breakdown of the association between mean transfer rate and TPJ connectivity during trust relative to control decisions in the presence of threat via the interaction contrast (Trust_no threat_ > Trust_threat_) > (NS Control_no threat_ > NS Control_threat_). We found a significant effect in the right STS (64, -43, 6, p < 0.05, SV FWE-corrected, k = 7; Supplementary Table 4b, recall (see online methods) that the STS is the most ventral part of our TPJ mask). To characterize the interaction effect, we ran post hoc simple effects analyses that compare the relationship between transfer rates and TPJ functional connectivity as a function of threat in the trust game. Specifically, we examined whether aversive emotion caused significant changes in the relationship between TPJ connectivity (with its target regions) and mean trust levels in the trust game. This analysis showed that aversive emotion caused a general breakdown of the association between left TPJ connectivity and mean trust in the right posterior superior temporal sulcus (64, -43, 6, p < 0.05 SV FWE-corrected, k = 31; Supplementary Table 5c). In this region, therefore, there was a significantly positive relationship between left TPJ connectivity and mean trust levels during the no-threat condition (Figure 3c, green regression line) that vanished in the presence of threat (Figure 3c, red regression line). In contrast, during decisions in the non-social control task no voxels in any of our ROIs, as well as the superior temporal sulcus showed greater connectivity during no threat relative to threat, or the reverse contrast of threat vs. no threat, even at a very liberal threshold of p < 0.05, uncorrected.

These results suggest that connectivity between left TPJ and its target region in contralateral pSTS supports trust when distortionary aversive emotion is absent. However, in the presence of aversive emotion, the relationship between the connectivity pattern in the trust network and behavioral trust was suppressed. Thus, aversive emotion not only reduced average trust but also diminished specific relationships between the connectivity patterns in the TPJ network and behavioral indices of trust. Our results therefore suggest that the pSTS is a crucial neural node that mediates the breakdown of trust in the presence of threat.

### Aversive emotion alters activation patterns within choice-relevant domain-general neural circuitry

The previous analyses indicate that aversive emotion had distinct effects on neural processes devoted to trust decisions and that functional connectivity strength between TPJ and its targets was specifically related to behavioral trust. However, the pronounced emotional reactions to the threatening context also had an impact on non-social control decisions. We therefore addressed the question to what extent aversive emotion impacts general choice-related neural circuitry in both the trust and the non-social control game via an exploratory whole-brain analysis investigating the main effect of aversive emotion: (Trust_threat_ + NS Control_threat_) > (Trust_no threat_ + NS Control_no threat_). This identified a domain-general network of regions showing either suppression or enhancement in choice-related neural activity during both the trust and the non-social control task (Figure 4 and Supplementary Table 7). The suppression of neural activity in the threat condition (red time course, Figure 4) relative to the no threat condition (green time course, Figure 4) was observed in both the trust and non-social control task (Supplementary Figure 4) in bilateral posterior dlPFC (left: -62, -4, 18, k = 686 and right: 62, -6, 28, k = 515), left amygdala (-24, -15, -23, k = 226), posterior paracentral lobule (4, -36, 69, k = 309), and left vlPFC (-48, 41, -8, k = 281) and vmPFC (-10, 44, -8, k = 464; Supplementary Table 6a). Significant enhancement of activity during decision-making under aversive emotion (Supplementary Figure 5) was obtained in the cerebellum (-4, -46, -24, k = 549; Supplementary Table 6b). Together, these results identify a network of domain-general regions whose decision-related activity is significantly impacted by incidental aversive emotion. Notably, the regions identified by the main effect do not overlap with regions showing trust-specific effects (tested via conjunction analysis), underlining that the trust-specific effects of aversive emotion occur above and beyond domain-general effects on decision-making.

**Figure 4.**
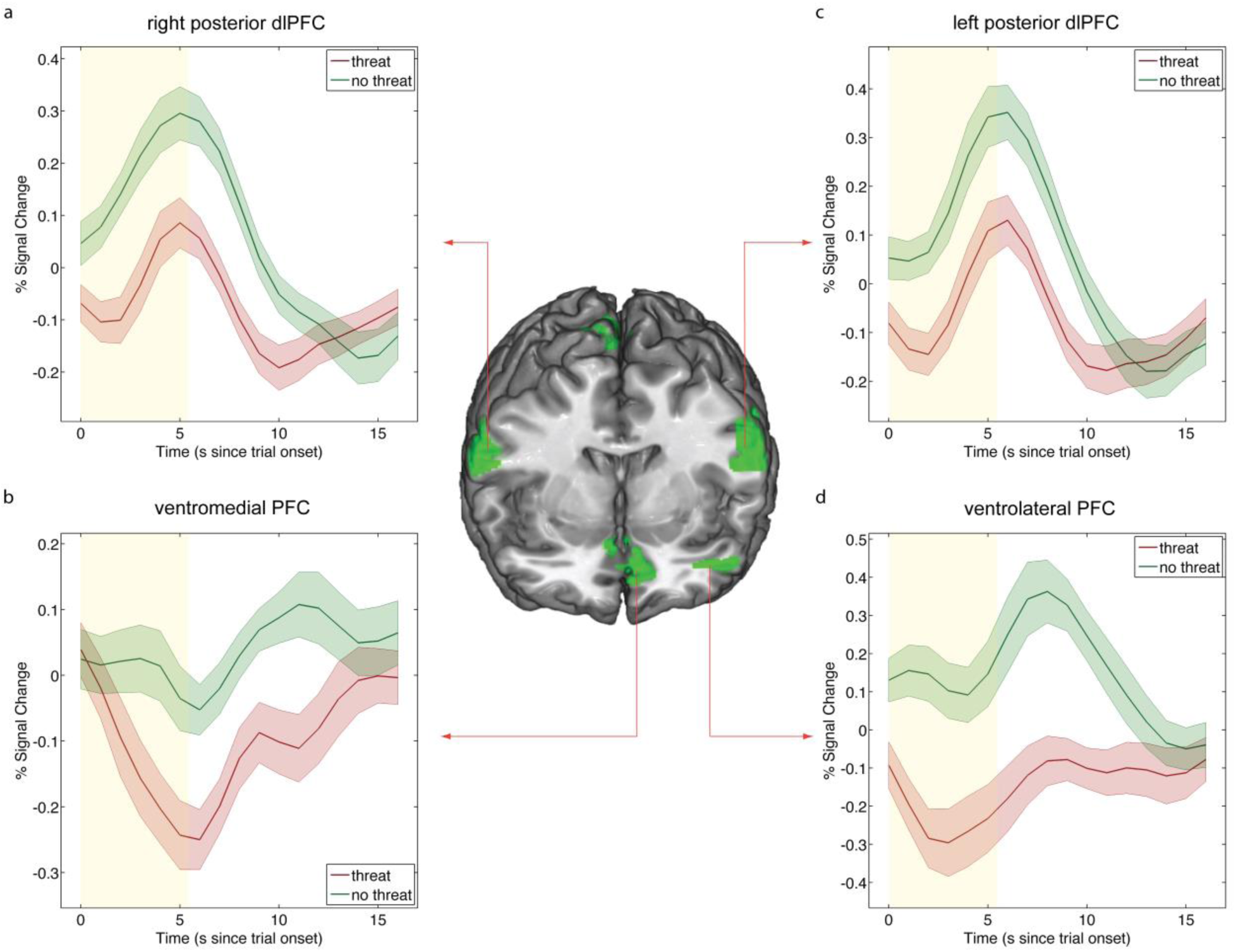
The impact of aversive emotion on choice-domain independent neural correlates of decision-making. We tested the main effect of aversive emotion on the neural correlates of decision-making independent of the choice domain (social and non-social). This analysis revealed a domain-general network consisting of bilateral posterior dlPFC (panel a, right peak at xyz = -62, -4, 18, and (panel c, left peak at xyz = 62, -6, 28), and two clusters in ventromedial prefrontal cortex (left peak at xyz = -10, 44, -8; right peak at xyz = 6, 21, -14 panel b) and left ventrolateral PFC (panel d, peak at xyz = -48, 41, -8). These regions show significant threat-related suppression (no threat > threat, regions shown in green) in choice-related activity during both trust and non-social control trials. Additional regions that are shown in Supplemantary Figure 4 include left amygdala (panel c, peak at xyz = -24, -15, -23), posterior paracentral lobule (peak at xyz = 4, -36, 69). Time courses reflect choice-domain independent activity that shows suppressions due to the aversive emotion during decisions in both trust and non-social control trials. To illustrate the equivalent effect of aversive emotion, Supplementary Figures 4 and 5 show activity for both trust and control trials in separate graphs. Time courses were extracted from 6 mm spheres around peak voxels. The 5.5-second choice period is displayed in yellow.

## Discussion

Incidental aversive emotion is a ubiquitous phenomenon that pervades many aspects of human behavior and human social interaction. In this paper, we investigated the behavioral and neural impact of incidental emotion on trust decisions. We employed a novel experimental technique to establish aversive emotion by inducing a prolonged expectation of unpredictable and aversive tactile stimulation embedded within a hybrid fMRI design. The threat of painful tactile stimulation significantly increased autonomic arousal during both social and non-social decision-making and was associated with consistent self-reports of the experience of aversive emotion. We observed that aversive emotion significantly reduced subjects’ trust in their partners. To the extent to which aversive emotions are associated with stress, this result is consistent with a recent behavioral study that showed that acute stress reduces trust ^6^. Importantly, despite the fact that aversive emotion was incidental to the decisions made by subjects, we observed significant behavioral and neural effects of the aversive emotional state. This finding contradicts consequentialist economic models that assume that emotions can at most affect choices by changing subjects’ preferences over outcomes^12^. Incidental emotions, which by definition are unrelated to the consequences of ongoing decisions, are therefore assumed to exert no influence on decision-making. Our behavioral results underline the importance of emotions in decision-making, even when they are unrelated to choice outcomes and clearly contradict the assumption of consequentialist economic models. Our neuroimaging results highlight mechanisms by which incidental emotions can impact on social decision-making, namely by suppressing the social cognitive neural processes that enable predictions about the interaction partner’s intentions.

Our neuroimaging results provide information about the neural mechanisms behind the reduction of participants’ trust. We show a disruption of trust-specific neural activity and connectivity due to aversive emotion. While the temporoparietal junction (TPJ) was preferentially engaged during trust decisions, aversive emotion led to a trust-specific breakdown of this activation pattern (Figures 2a-b) in the left TPJ (and in right TPJ at a reduced threshold). Moreover, in the absence of aversive emotion, we observed significant connectivity between TPJ and amygdala during trust decisions, but this connectivity was disrupted when we induced aversive emotion (Figures 2c-d). Aversive emotion also disrupted the relation between the connectivity patterns in the neural network underlying trust and the magnitude of behavioral trust. In particular, functional connectivity strength with the TPJ predicted mean trust levels for a network of regions consisting of amygdala, DMPFC, superior temporal sulcus (STS), and VLPFC (Supplementary Tables 5 and 6, Figures 3a-b, d). Aversive emotion caused a specific breakdown of this association for the posterior STS, such that the connectivity between left TPJ and right pSTS no longer predicted overall trust-taking (Figure 3c). Our results therefore identify a network of interconnected regions consisting of left TPJ, amygdala and right pSTS, for which connectivity patterns during trust-taking are significantly impacted by incidental aversive emotion.

The previous literature ^3,28,61^ has identified betrayal aversion as one of the key determinants of trust decisions in the trust game. Betrayal aversion means that subjects find it extremely aversive to be cheated in the trust game by an untrustworthy partner. Betrayal aversion therefore constitutes a powerful motivation to form expectations about the partner’s responses to the various trust levels and to assess the emotional significance of these responses. These processes critically involve subjects’ mentalizing faculties and the assessment of the (negative) emotional value of the possibility of being cheated by an untrustworthy partner ^23,25,27,32,34,62^.

The temporoparietal junction (TPJ) and the DMPFC have repeatedly been implicated in representing and interpreting others’ mental states and behavior ^35,36,47,63-65^ and the amygdala has been shown to respond strongly to variations in the trustworthiness of faces ^54,55^ and other emotionally salient stimuli ^66,67^. We therefore conjectured that these regions also play a key role in the computations involved in assessing and evaluation the partner’s anticipated responses in the trust game. This hypothesis is consistent with prior reports of TPJ, DMPFC and amydgala activation during trust decisions ^23,25,32,34,68,69^. Our present results significantly extend these prior findings, by underlining the behavioral relevance of interacting neural networks (rather than just isolated areas) for trust decisions. This is most consistently shown by our findings that enhanced connectivity between TPJ and regions important for social cognition (DMPFC, STS) and emotion processing (amygdala, VLPFC) relate to each individuals’ levels of trust.

Most importantly, however, we show that key components of these trust-supportive and trust-specific neural networks are suppressed by aversive emotion. In particular, we find that threat of shock leads to (1) specific reductions of TPJ activation during trust decisions; (2) specific reductions in the connectivity between TPJ and amygdala during trust decisions; (3) general reductions in choice-related activity in the amygdala; and (4) trust-specific disruptions of the association between TPJ connectivity and mean trust. These results thus suggest that aversive emotions undermine decisive components of trust-specific neural networks involved in the computations relevant for assessing a partner’s responses to various trust levels and the associated emotional valuations. This effect is expressed particularly clearly in the change in TPJ-amygdala connectivity due to threat-of-shock: In the absence of threat, TPJ and amygdala show trust-specific communication likely reflecting the integration of social cognitive (mentalizing, TPJ) and social emotional (trustworthiness assessments and evaluations, amygdala) information important for trust decisions. In the presence of threat, by contrast, the amygdala shows a general suppression due to the emotional context (Supplementary Figure 4) and this preoccupation with the immediate and emotionally highly salient threatening context seems to prevent it from communicating with TPJ. In other words, aversive emotions seem to reduce a subject’s capacity to mentalize about and evaluate the emotional consequences of various trust levels and, as a consequence, subjects reduce their behavioral trust towards their partner.

It is important to note that at the behavioral level, we observed that aversive emotions reduced both trust and non-social control decisions. This result is not surprising, given that the fMRI design was conceptualized and optimized to reveal the effects of aversive emotion on the neural correlates of *trust decisions*, which required the non-social control task to be equivalent to the trust game in all respects, except that a randomization procedure and not another human being decided the backtransfer amount. This finding therefore underlines that the induction of aversive emotion via threat of shock was successful. The comparable impact of aversive emotion on trust and nonsocial control trials furthermore confirms that our non-social condition constitutes a well-matched control for the trust game. Importantly, we show that despite the similar impact of aversive emotion on trust and control decisions, subjects performed differently in these two game types, as evidenced by (1) significant differences in average transfer rates and reaction times, (2) similarly strong correlations within a game type, but significantly different correlations across game types (but within shock conditions) and (3) an absence of a significant correlation between the threat-induced differences in transfer rates in the trust and control games. These three sources of evidence underline that different mental processes underlie decisions across the trust and non-social control games and that aversive emotion therefore impacted separable underlying cognitive processes. These results are consistent with previous observations that two experimental manipulations can generate behavioral effects of similar size, but still reflect influences on very different mental operations (e.g., trust vs. general risk-taking) as evident by different brain activity ^70-75^. Moreover, if the purpose of the study is to understand these differences in terms of brain activity and connectivity, it is in fact desirable that there are no differences in behavioral responses across conditions (e.g., ^70-74^; see also ^75^).

An alternative explanation for our observation of suppression in behavioral trust and its neural correlates in TPJ is that subjects may have simply been distracted by the presence of threat. However, our finding of significantly faster response times in the presence of threat makes this alternative interpretation unlikely, as this pattern of results is opposite to what would be expected if participants were distracted or under heavier cognitive load. Numerous studies have consistently shown that a hallmark of cognitive load (or distraction) is a slowing of response times, and, in fact, longer RTs are often used as a validation that an experimental load or distraction manipulation worked ^76^. Our findings that response times were faster under threat is therefore much more consistent with the interpretation that participants were emotionally aroused under threat ^77^ and that this emotional arousal impacted social decisions and their neural correlates.

In conclusion, we report results that show a significant behavioral impact of incidental emotions on trust-taking and we identify the trust-specific neural mechanisms associated with the impact of aversive emotion on trust. These effects were observed even though induced emotions were unrelated to the choice outcomes in our task, confirming that incidental emotions can have a powerful impact on behavior and its underlying mental operations. Our findings inform the development of economic and social theory and call for the integration of incidental emotion in behavioral models of social and non-social decision-making ^12-14^. In addition, by identifying the specific distortions of the neural network activity supporting trust, we provide a first step towards neural models that help us better understand such distortions. In particular, our results support the notion that an important mechanism through which aversive incidental emotion impacts social decision-making is the suppression of activity and connectivity between regions known to be crucial for mentalizing about other people’s responses (such as the temporoparietal junction, dorsomedial PFC and the superior temporal sulcus) and the evaluation of socially threatening stimuli (such as the amygdala). Given that psychiatric diseases, such as pathological anxiety, social phobia or depression, are characterized by a particularly pronounced susceptibility to negative emotion, our results may also be useful in understanding the neural circuitry associated with emotion-related distortions of social behavior in psychiatric diseases.

## Methods

### Participants

41 human volunteers (mean age (std.) = 22 (2.145), 17 females) from various Universities in Zurich participated in the current experiment. Only right-handed subjects between the ages of 18 – 45 with no prior psychiatric illness, no regular illicit drug use and no traumatic head injury were included in the experiment. The sample size was based on recommendations for investigations of individual difference in fMRI studies based on power simulation ^78^. All participants gave written informed consent to procedures approved by the local ethics committee (Kantonale Ethikkommission, Zurich, Switzerland) before participating in the study. Subjects were right-handed as assessed by the Edinburgh handedness questionnaire and did not report any history of psychological illness or neurological disorders, as assessed by a standard screening form.

### Prescanning procedure

Particular care was taken to ensure that subjects understood all aspects of the experiment. To this end, subjects were instructed to carefully read detailed instructions and were required to fill out an extensive questionnaire probing their understanding of the experimental procedures. The accuracy of each subject’s answers was confirmed by the experimenters and discussed in a brief interview that lasted for ca. 10-minutes. Subjects were then placed inside the scanner for a brief practice session consisting of 12 trials to ensure that they could view all stimuli, perform the task, make decisions in the allotted 5.5 seconds per trial, understood the experimental setup and to give subjects the opportunity to ask further questions.

After completion of practice, subjects were taken out of the scanner and washed their hands before placement of SCR and stimulation electrodes. Subjects were then placed inside the scanner and two ring electrodes were attached to the dorsum of the left hand: (1) the electrode providing relatively higher intensity stimulation was placed between one to two cm below the second carpometacarpal joint, and (2) the electrode providing relatively lower intensity stimulation was placed one to two cm below the fifth carpometacarpal joint. To determine individual thresholds for high-intensity and low-intensity stimulation, we followed a standard procedure ^79,80^ and employed a visual analog rating scale (VAS) with endpoints defined as 0 = ‘cannot feel anything’ and 10 = ‘maximum tolerable pain’. Tactile stimulation was delivered via two Digitimer DS5 isolated bipolar constant current stimulators (bipolar constant current, 5V, 50mA, Digitimer Ltd, Welwyn Garden City, UK) and a custom-made fMRI compatible 5-mm ring electrode, which delivered a maximally focused and centered tactile stimulus. By varying current amplitude between 1 and 99 % of maximum amperage, stimuli with varying intensity levels were repeatedly delivered to each participant until stable ratings were achieved at least three times according to the following criteria: between 1 and 2 for the low intensity stimulus, and between 8 and 9 for the high intensity stimulus. Visual and tactile stimulus presentation, as well as recording of responses, were controlled by Cogent2000 (http://www.vislab.ucl.ac.uk/cogent.php).

### Task

To investigate the effect of incidental emotion on trust-taking, we employed a hybrid fMRI design, in which aversive emotion was manipulated in a blocked fashion while social (trust) and non-social (control) tasks were presented in an event-related fashion. Specifically, we varied aversive emotion by creating an expectancy of weak or strong unpredictable electrical stimulation that could occur at any time for the duration of an entire block. This expectancy was created by means of a block cue presented at the beginning of each block that informed participants about the game type (trust or control game) and the intensity of stimulation (weak or strong) for the current block (Figure 1a). Stimulation intensity was communicated to subjects in three ways: (1) via a verbal cue embedded in the 750-ms block cue “strong” for treatment (“threat” condition), “weak” for control (“no-threat” condition); (2) via a predictable tactile reminder cue presented 700 ms after visual cue onset that reflected the exact stimulation intensity of the current block; (3) via a specific background color that was consistently associated with either threat or no-threat blocks for each subject (color was counterbalanced across subjects) and remained constant for the duration of a block. The number and time points of electrical stimulation events throughout the blocks were determined to be completely unpredictable to subjects, in order to augment the efficacy of the threat-of-shock treatment. For this purpose, the number of stimulation events was determined for each block by random draw from a gamma distribution (shape parameter = 1; scale parameter = 1). Participants therefore experienced exactly one predictably reminder and, on average, one additional unpredictable electrical stimulation per block. The exact timing of these stimulation events was then determined at random time points between the offset of the cue display and onset of the resting screen drawing from a uniform distribution, with the constraint that at least 0.2 s separated successive electrical shocks. Timing and order of stimuli were randomized for each subject to maximize identification of the effects of aversive emotion on the neural correlates of trust decisions using in-house software programmed in Matlab.

Each block commenced with the set of cues described above that indicated the type of decision to be made (non-social control or trust) and the level of stimulation (weak, strong) to be expected by subjects for the rest of the block. After a brief and jittered interstimulus interval of 3-9 seconds, the first of three trials within a given block was displayed. In both the trust and the control game, subjects were presented with a multiple-choice scenario, in which one of five amounts between 0 and 24 Swiss Francs (CHF) could be transferred to Player B or invested in a lottery. While subjects always had the options to either invest all (24 CHF) or none (0 CHF) of their endowment, each trial presented a novel choice scenario by (1) varying the intermediate options between 4, 6, or 8 CHF in the low category, 10, 12, or 14 CHF in the medium category, and 16, 18, or 20 in the high category of intermediate transfer amounts; (2) varying the location of each choice option and (3) varying the location of the originally highlighted choice option. This variability was introduced in order to ensure that subjects paid attention to all choice options on every trial and to avoid excessive use of heuristics. Intermediate amounts, location of choice options and location of the initially highlighted choice option were fully counterbalanced across conditions. Subjects selected their preferred option by moving a yellow dot that highlighted the currently selected choice option up and down by pressing two dedicated buttons on a standard MR-compatible 4-button response box and confirming their choice by pressing a third button. At this point the selected choice option was highlighted in red for the remaining duration of a trial. After a jittered intertrial interval (3-9 seconds) a new trial began. Please note that in order to control for wealth effects, subjects in our experiment did not receive trial-by-trial feedback about the financial outcome of their choices in both the trust and the non-social control game. By using one-shot games with no feedback we preclude learning- and outcome-related signals commonly observed in valuation regions (e.g., ^63,81^). Subjects completed 28 blocks (7 blocks per condition with an average length of 38.75 seconds) with three trials each in two runs.

### Payment determination

We collected trustee responses in separate behavioral sessions that were conducted prior to the fMRI experiment using the same trust game. We elicited the trustees’ choices with the strategy method, i.e. the trustees indicated their responses to each feasible transfer level. The trustees gave written and informed consent that we could use their strategies in follow-up experiments. In the fMRI part of our experiment the subjects (investors) thus played against the pre-recorded strategies of the trustees, i.e. a subject’s transfer level together with the strategy of the (randomly) matched trustee determined the final monetary outcome in a trust game trial. Given the absence of the trustee on the scanning day, we informed participants that they were interacting with trustees in a temporally delayed fashion. Specifically, we emphasized to subjects that their payoffs were determined by decisions of real persons in the trust game, and by a computer algorithm in the control game, and that they were assigned different real persons across trust game trials. Finally, to maintain the interactive nature of the trust game, we informed our subjects that their choices had real, but delayed, consequences for trustees, who were sent additional payments according to the decisions made by the investor in the scanner after completion of the experiment. During the experiment, the subjects did not receive any feedback about the behavior of their matched trustees, or the payoff amounts from lottery investments.

After completion of the fMRI part of the experiment, the subjects selected two trials at random by dice throw and payment was determined according to the decisions made by the subject and the trustee on the selected trials. In order to avoid hedging, both payout trials were drawn from the entire experiment, i.e. the payout trials were not specific to a condition, such as the trust or control game. If a trust game was randomly chosen for payout determination, the investor’s payout was determined based on the amount transferred to the trustee and the backtransfer amount of the specific trustee the investor was paired with on that trial (payout investor = 24 – transfer to trustee + backtransfer from trustee; payout trustee = 24 + transfer from investor * 3 – backtransfer to investor). If a control game was randomly chosen, the computer algorithm randomly drew a payout amount from the distribution of trustees’ backtransfer amounts. Our procedure therefore created equivalent payout amounts and likelihoods for the trust and control game.

### Exit questionnaire

After completion of the experiment, subjects filled out an exit questionnaire that probed their beliefs about the accuracy of our instructions, as well as emotional reactions to our experimental manipulations. The main goal of the exit questionnaire was to measure whether subjects believed our instructions. Note that we implement such measurements routinely although we have little reason to believe that subjects doubt our instructions. Our laboratory uses deception only as a very rare exception, and we also did not use any deception in this experiment and fully disclosed all information truthfully to the subjects. Subjects were asked to rate 7 statements on a scale from 0, indicating very unbelievable, to 4, indicating very believable. The statements declared that the trust games were played with real persons, that each trust game was played with an anonymous trustee, that decisions of trustees were made by actual persons, and that trustees will receive additional payments based on the decisions of subjects on the relevant trial. Subjects’ responses were entered into one-sample t-tests testing whether responses were significantly greater than the mid-point of the scale (2, indicating neither believable nor unbelievable). Mean ratings for all statements were significantly greater than two, indicating that subjects believed the statements (all t-tests survive the Bonferroni-corrected threshold of 0.007; average rating (SD) over all statements is 3.37 (0.86)).

### Skin conductance responses (SCRs)

Skin conductance responses were collected using a PowerLab 4/25T amplifier with a GSR Amp (ML116) unit and a pair of MR-compatible finger electrodes (MLT117F), which were attached to the participants’ left middle and ring finger via dedicated Velcro straps after application of conductance gel. Subjects’ hands had been washed using soap without detergents before the experiment. Stable recordings were ensured before starting the main experiment by waiting for signal stabilization during training and stimulation intensity calibration. LabChart (v. 5.5) software was used for recordings, with the recording range set to 40 µS and using initial baseline correction (“subject zeroing”) to subtract the participant’s absolute level of electrodermal activity from all recordings (all specs for devices and electrodes from ADInstruments Inc., Sydney, Australia).

Due to technical problems, data from 4 subjects included 1 run (out of 2) and data from 1 subject was lost. Each participant’s SCR data were initially smoothed with a running average over 500 samples (equivalent to 500 ms at a sampling rate of 1KHz) to reduce scanner-induced noise. Data were then resampled from 1KHz to 1Hz and subsequently z-transformed. Statistical analysis of the pre-processed skin conductance data followed the approach commonly employed in analyses of fMRI data. Specifically, multiple linear regression implemented in AFNI was used to estimate SCR during decisions made in each of the task conditions, that is, during trust and non-social control tasks and in the context of threat and no-threat treatment blocks. The statistical model included a total of 7 regressors that reflected the onset times of decision screens in trust and non-social control trials under expectancy of strong and weak electrical shocks, cue times indicating the onset of a block, as well as delivery times of strong and weak tactile stimulation. To avoid making assumptions about the shape of the SCR response, a finite impulse response (FIR) model was used to estimate average responses (beta weights) during each trial type via deconvolution from event onset to 16s post onset using 17 cubic spline basis functions. Constant, linear and quadratic terms were included as regressors of no interest for each run separately to model baseline drifts of the SCR. Regressor estimates (beta weights) at each time point and for each condition were then used in follow-up analyses reported in the Results section.

### fMRI data acquisition

Magnetic resonance images were collected using a 3T Philips Intera whole-body magnetic resonance scanner (Philips Medical Systems, Best, The Netherlands) equipped with an 8-channel Philips sensitivity-encoded (SENSE) head coil. Structural image acquisition consisted of 180 T1-weighted transversal images (0.75-mm slice thickness). For functional imaging, a total of 1095 volumes were obtained using a SENSE T2*-weighted echo-planar imaging sequence ^82^ with an acceleration factor of 2.0. We acquired 45 axial slices covering the whole brain with a slice thickness of 2.8 mm (inter-slice gap of 0.8 mm, sequential acquisition, repetition time = 2470 ms, echo time = 30 ms, flip angle = 82°, field of view = 192 mm, matrix size = 68 × 68). To optimize functional sensitivity in orbitofrontal cortex and medial temporal lobes, we used a tilted acquisition in an oblique orientation at 15° relative to the AC-PC line.

### fMRI data analysis

Preprocessing and statistical analyses were performed using SPM8 (Wellcome Department of Imaging Neuroscience, London, UK). To correct for head motion, all functional volumes were realigned to the first volume using septic b-spline interpolation and subsequently unwarped to remove residual movement-related variance due to susceptibility-by-movement interactions. Slice timing correction was performed after realignment/unwarping. To improve coregistration, bias-corrected anatomical and mean EPI images were created and subsequently coregistered using the new segment toolbox in SPM. Images were normalized to the Montreal Neurological Institute T1 template using the parameters (forward deformation fields) derived from the nonlinear normalization of individual gray matter tissue probability maps. Finally, functional data underwent spatial smoothing using an isotropic 6-mm FWHM Gaussian kernel.

Statistical analyses were carried out using the general linear model. Regressors of interest were modeled using a canonical hemodynamic response function (HRF) with time and dispersion derivatives in order to account for subject-to-subject and voxel-to-voxel variation in response peak and dispersion ^83^. Since our main interest was the impact of aversive emotion on trust-taking, we modeled the decision period for the full response time on each trial, that is from the onset of the decision screen until subjects pressed the confirm button. This was done in the following four conditions: (1) trust game during relatively high-intensity stimulation expectancy (threat condition), (2) trust game during relatively low-intensity stimulation expectancy (no-threat condition), (3) control game during relatively high-intensity stimulation expectancy (threat condition) and (4) control game during relatively low-intensity stimulation expectancy (no-threat condition). Finally, the following regressors of no interest were included in our model: the actually realized weak and strong tactile stimulations during a block (one reminder shock and, on average, one additional shock randomly drawn from a gamma distribution were administered per block), block cues indicating game type (trust, control) and stimulation intensity of the reminder shock (weak, strong) at the beginning of each block, as well as omissions of behavioral responses during a trial.

The main goal of the current investigation was to identify the impact of aversive emotion on the neural correlates of trust decisions. Trust-specific neural effects of aversive emotion can be identified via an interaction between threat and game type, in which threat significantly alters the neural correlates of decision-making in trust relative to non-social control trials. To investigate the interaction between threat and game type, an ANOVA was computed by entering contrast estimates obtained from first level models into a flexible factorial model with the factors game type (trust, control), threat (absent, present), as well as subject. We were particularly interested in trust-specific emotion-induced suppression of activity and connectivity, which we tested via the interaction contrast (Trust_no threat_ > Trust_threat_) > (NS Control_no threat_ > NS Control_threat_) in the context of the flexible factorial design. A covariate reflective of each subject’s mean transfer in each condition was also included to probe for brain-behavior correlations. All analyses were also conducted without the behavioral covariate and results did not change.

Our main analyses rely on a priori regions of interest as we expected regions commonly implicated in the major cognitive and affective component processes of trust to be affected by aversive emotion. Specifically, we hypothesized that subjects needed to assess the trustworthiness of the trustee to make predictions about payout probability in the trust game, which involves regions commonly implicated in theory of mind and social cognition ^84,85^. To identify regions implicated in theory of mind, we consulted neurosynth.org ^57^, which offers a means to obtain automated meta-analyses over a large number of prior fMRI investigations and thereby provides an independent method to obtain masks for ROI analyses. To guide and constrain our ROI selection, we computed the conjunction of the neurosynth meta-analyses for the terms “emotion” (forward inference to identify regions that *consistently* show modified activity) and “theory of mind” (reverse inference to identify regions that are *specifically* involved in theory of mind). This approach identified overlap between these networks in left TPJ and DMPFC, which agrees particularly well with results from recent a meta-analysis identifying the TPJ and DMPFC as core social cognition regions^5^. Furthermore, because of its prominent role in signaling emotional salience ^15^, we included the amygdala as an additional ROI (see also neurosynth search: “emotion”).

ROI analyses in relevant cortical regions were conducted using small volume correction with masks created via relevant search terms on neurosynth.org, while anatomically well-defined subcortical ROI masks were created using the AAL atlas implemented in WFU Pickatlas. The following independent ROI masks were created via automated meta-analyses from neurosynth.org: (1) bilateral temporoparietal junction (neurosynth term: theory of mind), with peak voxels in left (-60,-56,14) and right TPJ (56,-58,20) and sizes of 1031 and 1416 voxels, respectively. It is noteworthy that the ventral part of this mask also contains voxels from posterior superior temporal sulcus (STS). For simplicity, we refer to this mask nevertheless as “TPJ”. (2) dorsomedial PFC (neurosynth term: theory of mind), with a peak voxel in medial DMPFC (-2, 28, 62) and a size of 3175 voxels. The ROI mask for the amygdala, which is an anatomically well-defined region, was created via: bilateral amygdala (AAL) with sizes of 439 (left) and 492 (right) voxels. Additional exploratory analyses were conducted in regions involved in evaluating the anticipated outcomes of choice options, such as ventral striatum and vmPFC ^86^ (neurosynth term: “reward”), as well as a region implicated recently in one-shot trust decisions by a recent meta-analysis ^51^ using the following masks: bilateral ventral striatum (combined mask of AAL putamen and caudate up to z = 8), with sizes of 3239 (left) and 3429 (right) voxels, respectively; ventromedial PFC (neurosynth search term: ventromedial) with a peak voxel in medial vmPFC (8, 24, -12), with a size of 1327 voxels; left anterior insula with peak voxels at -42, 18, 2 and a size of 13844 voxels and right anterior insula with peak voxels at 42, 18, 2 and a size of 13871 voxels.

Furthermore, to identify whether extended networks outside our regions of interest show effects of interest, we conducted exploratory whole brain analyses at an FWE-corrected extent threshold of p < 0.05 (k > 226, initial cluster-forming height threshold p < 0.001). Finally, to characterize activation patterns of interest, such as time courses and activation differences due to aversive emotion, regression coefficients (beta weights) for the canonical HRF regressors were extracted with rfxplot ^87^ from 6 mm spheres around individual subjects’ peak voxel that showed significant effects of interest on BOLD responses and functional connectivity. Follow-up tests that characterize the single components of significant interaction effects were conducted in neuroimaging space via tests of simple effects of interest.

### PPI analyses

Psychophysiological Interaction analyses were conducted using the generalized form of context-dependent psychophysiological interactions toolbox (gPPI toolbox ^60^), using the same statistical model as outlined above. All voxels that survived SV FWE-correction for the interaction contrast in left TPJ (-60, -54, 19, k=95) were used as seed region (shown in blue color in Figure 2c). To obtain an estimate of neural activity within the seed region, the BOLD signal from the seed region was extracted, corrected by removing effects of noise covariates, and deconvolved ^88^. Psychological interaction regressors for each of the task type and stimulation intensity combinations control decisions during (1) weak and (2) strong stimulation, trust decisions during (3) weak and (4) strong stimulation were created by multiplying the estimated neural activity during the relevant decisions with condition-specific on- and offset times convolved with the canonical HRF. A new GLM was then estimated for each subject that consisted of the original design matrix with the addition of the four psychological interaction regressors and the time course from the seed region.

To investigate the impact of aversive emotion on trust-specific functional connectivity of the left TPJ, we probed the functional connectivity data for an interaction between threat and game type. To investigate the interaction between threat and game type, we entered the contrast estimates obtained from first level PPI models into a flexible factorial model with the factors game type (trust, non-social control), threat (absent, present), and separate covariates reflecting mean transfer in each condition. A subject factor was also included in the model. Given that we were particularly interested in trust-specific changes in functional connectivity, we first contrasted the covariates reflecting mean transfers in the trust game and mean transfers in the non-social control task in the absence of threat (Trust_no threat_ > NS Control_no threat_). This comparison identifies regions for which connectivity with the TPJ correlates more strongly with mean transfers in the trust game than with mean transfers in the non-social control game. As the next step, we then examined how threat of shock changed the relationship between TPJ connectivity and mean transfers in the trust game, by examining the interaction between game type and threat estimated over the covariates. We illustrate these results in Figure 3, by regression plots generated with coefficients reflecting functional connectivity strength for each of the conditions, extracted from 6-mm spheres around the peak voxel of the interaction contrast. The displayed regression plots were generated by the following regression model implemented in R (using the Regression Modeling Strategies package, RMS):

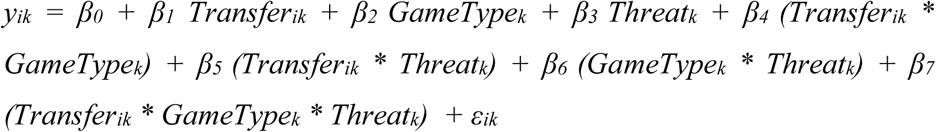

The dependent variable *y_ik_* is the functional connectivity strength between a given brain region and the TPJ for individual *i* in Game type k. Transfer_*ik*_ is the mean amount sent by individual *i* in Game type k. Game type is a dummy variable encoding whether decisions were made in the trust or the control task (1 indicates trust, 0 indicates non-social control task). Threat is a dummy variable encoding whether decisions were made in the presence or absence of threat (1 indicates presence, 0 indicates absence of threat). In this regression, the coefficient for Transfer_*ik*_ (β_*1*_) measures the slope of the relationship between TPJ connectivity and mean transfers in the absence of threat in the control task (see blue lines in Figures 3a – 3c), and the sum of the coefficients for Transfer_*ik*_ (β_*1*_) and the interaction term between Transfer_*ik*_ * GameType_*k*_ (β_*4*_) measures the trust-related slope increase of the relationship between TPJ connectivity and mean transfers in the absence of threat (see orange lines in Figures 3a – 3c). Equivalent analyses were performed to probe for significant differences between the threat and the no-threat condition in the trust game in the relationship between functional TPJ connectivity and mean transfer levels. Here, the sum of the coefficients for Transfer_*ik*_ (β_*1*_), the interaction term between Transfer_*ik*_ * GameType_*k*_ (β_*4*_), the interaction term between Transfer_*ik*_ * *Threatk* (β_*5*_), and the interaction term between Transfer_*ik*_ * GameType_*k*_ * Threat_*k*_ (β_*7*_) measures the slope of the relationship between TPJ connectivity and mean trust in the presence of threat (see red line in Figures 3d).

## Supplementary Information

### 1 Supplementary Analyses

#### 1.1 OLS regression investigating the influence of experienced electrical stimulation on choice

It is possible that the main driving force of the behavioral change that we report in the main paper is not due to the affective impact of our threat of shock manipulation, but to the actual experience of electrical stimulation immediately prior to decision-making. To assess this possibility, we investigated the effects of experienced electrical stimulation on choice behavior in the trust and the non-social control game. To this end, we ran comprehensive ordinary least-squares (OLS) regression analyses in R version 2.5.12. These analyses predicted for each individual *i* the observed choice T_i,t_ on trial *t* with the following equation:

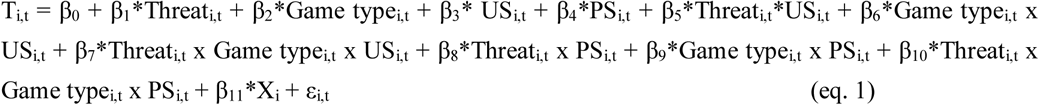

T_i,t_ reflects the transfer amounts to either a lottery or Player B on a given trial. *Threat*_*i,t*_ is an effect-coded variable (1 = threat of shock, -1 = safety) reflecting the *expectation* of impending painful electric shock during the current choice scenario, not the *experience* of shock. *Game type*_*i,t*_ is an effect-coded variable (1 = trust game, -1 = control game) reflecting transfers within the context of a trust game. US_i,t_ and PS_i,t_ are effect-coded variables (1 = shock, -1 = no shock) encoding the *experience* of at least one unpredictable (US) and, respectively, predictable shock (PS) in the interval from 5 seconds prior to the display of the choice scenario until button press. We included PS in our regression models, in order to control for the influence of reminder shocks presented during the block cue. Of note, we also investigated shorter and longer intervals from 1 to 10 seconds prior to the choice scenario in 1-second increments and in all cases only *Threat* and *Game type* reach statistical significance, no other factors or interactions. The model also contains a constant parameter (β_0_), which measures the average transfer to the lottery in the no threat condition. Finally, the model includes a set of mean-centered socio-economic control variables (*X*_*i*_, e.g., age and gender) and relevant interaction terms that reflect differential effects of experiencing shocks on transfer behavior in different Game type and emotion contexts. We employed a random-effects model with robust standard errors adjusted for clustering on the subject level.

The results show significant treatment (threat) and game type (trust) effects, which we also report in the ANOVA analyses in the main paper. At the same time, regression results do not show significant effects of both unpredictable (p = 0.23) and predictable (p = 0.62) *experienced* electrical stimulation on choice behavior. Specifically, we do not observe a significant effect of, or interaction with, both unpredictable and predictable electrical stimulation. Together, regression results support and extend the results reported in the main paper. Specifically, threat of shock remains a significant predictor of choice behavior (p < 0.001) in the trust and the non-social control game, even when controlling for the presence of actually experienced electrical stimulation. Furthermore, we show that the experience of shock does not significantly impact behavior in any of our analyses, indicating that the *expectancy* of shock, not the *experience* of shock, significantly impacted choice behavior.

**Supplementary Table 1.** OLS Regression results reflecting the influence of experienced electrical stimulation on choice behavior

| Dependent Variable: Amount Transferred |  |  |  |  |  |  |
| --- | --- | --- | --- | --- | --- | --- |
|  | (1) |  | (2) |  | (3) |  |
| Threat | -0.659 | **** | -0.5482 | **** | -0.627 | **** |
|  | (0.1711) |  | (0.1389) |  | (0.1596) |  |
| Game Type | 1.2479 | **** | 1.2468 | **** | 1.338 | **** |
|  | (0.3242) |  | (0.3094) |  | (0.3097) |  |
| Game Type * Threat | -0.0128 |  | 0.0071 |  | -0.0654 |  |
|  | (0.1334) |  | (0.1107) |  | (0.1185) |  |
| Unpredictable S | 0.0146 |  | 0.0161 |  |  |  |
|  | (0.1179) |  | (0.1172) |  |  |  |
| Threat * Unpredictable S | -0.0649 |  | -0.0847 |  |  |  |
|  | (0.1254) |  | (0.1232) |  |  |  |
| Game Type * Unpredictable S | -0.1666 |  | -0.167 |  |  |  |
|  | (0.1170) |  | (0.1186) |  |  |  |
| Game Type * Threat * Unpredictable S | 0.1002 |  | 0.099 |  |  |  |
|  | (0.1413) |  | (0.1422) |  |  |  |
| Predictable S | 0.0011 |  |  |  | 0.0064 |  |
|  | (0.1237) |  |  |  | (0.1255) |  |
| Threat * Predictable S | -0.1837 |  |  |  | -0.1941 |  |
|  | (0.1099) |  |  |  | (0.1069) |  |
| Game Type * Predictable S | 0.0017 |  |  |  | -0.0198 |  |
|  | (0.1035) |  |  |  | (0.1061) |  |
| Game Type * Threat * Predictable S | -0.0303 |  |  |  | -0.0159 |  |
|  | (0.1023) |  |  |  | (0.1032) |  |
| Gender | 1.2645 |  | 1.2634 |  | 1.2659 |  |
|  | (1.3527) |  | (1.3533) |  | (1.3536) |  |
| Age | 0.264 |  | 0.264 |  | 0.2638 |  |
|  | (0.3253) |  | (0.3254) |  | (0.3258) |  |
| Intercept | 14.0029 | **** | 14.0041 | **** | 13.9996 | **** |
|  | (0.7103) |  | (0.7084) |  | (0.7103) |  |
| F | 17.26 |  | 24.78 |  | 24.72 |  |
| P | <0.001 |  | <0.001 |  | <0.001 |  |
| N | 3427 |  | 3427 |  | 3427 |  |
| AIC (corrected) | 22718.85 |  | 22712.35 |  | 22712.88 |  |

**Supplementary Table 2a.** ANOVA Results for Mean Transfer Rates

| Source | Type III Sum of Squares | df | Mean Square | F | Sig. | Partial Eta Squared |
| --- | --- | --- | --- | --- | --- | --- |
| Game Type | 299.438 | 1 | 299.438 | 20.319 | .000 | .337 |
| Error (Game Type) | 589.475 | 40 | 14.737 |  |  |  |
| Threat | 40.969 | 1 | 40.969 | 17.483 | .000 | .304 |
| Error (Threat) | 93.736 | 40 | 2.343 |  |  |  |
| Game Type * Threat | .179 | 1 | .179 | .122 | .729 | .003 |
| Error (Game Type * Threat) | 58.603 | 40 | 1.465 |  |  |  |

**Supplementary Table 2b.** ANOVA Results for Gender Effects on Mean Transfer Rates.

| Source | Type III Sum of Squares | df | Mean Square | F | Sig. | Partial Eta Squared |
| --- | --- | --- | --- | --- | --- | --- |
| Game Type | 302.822 | 1 | 302.822 | 20.180 | .000 | .341 |
| Game Type * Gender | 4.241 | 1 | 4.241 | .283 | .598 | .007 |
| Error(Game Type) | 585.235 | 39 | 15.006 |  |  |  |
| Threat | 50.086 | 1 | 50.086 | 26.623 | .000 | .406 |
| Threat * Gender | 20.365 | 1 | 20.365 | 10.825 | .002 | .217 |
| Error(Threat) | 73.371 | 39 | 1.881 |  |  |  |
| Game Type * Threat | .213 | 1 | .213 | .142 | .709 | .004 |
| Game Type * Threat * Gender | .069 | 1 | .069 | .046 | .831 | .001 |
| Error(Game Type*Threat) | 58.534 | 39 | 1.501 |  |  |  |

**Summary of gender effects on transfer amounts:** There was no main effect of gender $F(1,39) = 0.74$ , $p = 0.396$ , $\eta^2 = 0.019$ . The significant 2-way interaction between Threat and Gender is driven by females transferring significantly less on average during the Threat condition, independent of Game Type [Mean Transfer during Threat: 12.36 vs. Mean Transfer during no Threat: 14.197] compared to males [Mean Transfer during Threat: 14.35 vs. Mean Transfer during no Threat: 14.75].

**Supplementary Table 3a.** ANOVA Results for Mean Choice Latencies.

| Source | Type III Sum of Squares | df | Mean Square | F | Sig. | Partial Eta Squared |
| --- | --- | --- | --- | --- | --- | --- |
| Game Type | .411 | 1 | .411 | 6.226 | .017 | .135 |
| Error(Game Type) | 2.640 | 40 | .066 |  |  |  |
| Threat | .689 | 1 | .689 | 17.010 | .000 | .298 |
| Error(Threat) | 1.620 | 40 | .041 |  |  |  |
| Game Type * Threat | 3.396E-6 | 1 | 3.396E-6 | .000 | .993 | .000 |
| Error(Game Type*Threat) | 1.930 | 40 | .048 |  |  |  |

**Supplementary Table 3a.**
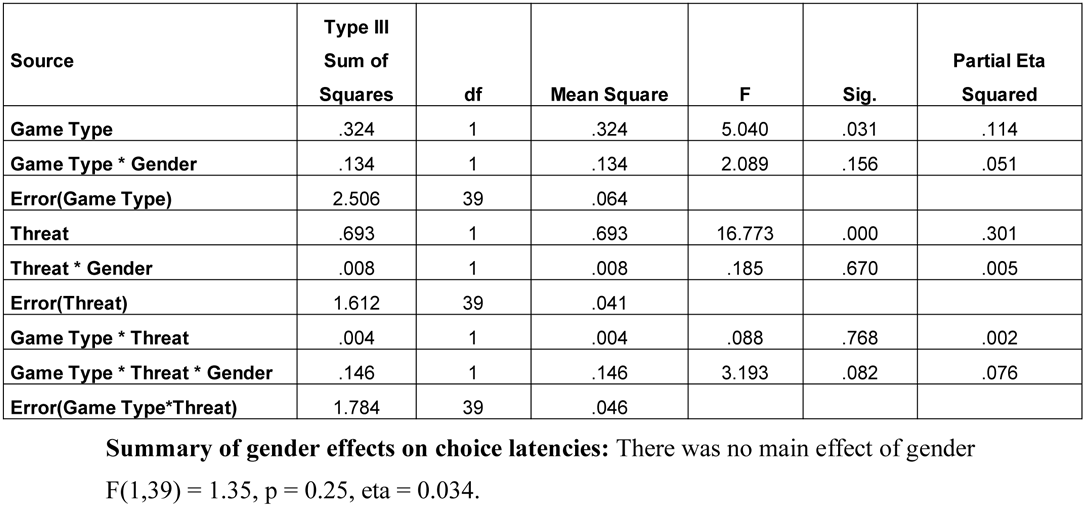
ANOVA Results for Gender Effects on Mean Choice Latencies.

| Source | Type III Sum of Squares | df | Mean Square | F | Sig. | Partial Eta Squared |
| --- | --- | --- | --- | --- | --- | --- |
| Game Type | .324 | 1 | .324 | 5.040 | .031 | .114 |
| Game Type * Gender | .134 | 1 | .134 | 2.089 | .156 | .051 |
| Error(Game Type) | 2.506 | 39 | .064 |  |  |  |
| Threat | .693 | 1 | .693 | 16.773 | .000 | .301 |
| Threat * Gender | .008 | 1 | .008 | .185 | .670 | .005 |
| Error(Threat) | 1.612 | 39 | .041 |  |  |  |
| Game Type * Threat | .004 | 1 | .004 | .088 | .768 | .002 |
| Game Type * Threat * Gender | .146 | 1 | .146 | 3.193 | .082 | .076 |
| Error(Game Type*Threat) | 1.784 | 39 | .046 |  |  |  |

**Summary of gender effects on choice latencies:** There was no main effect of gender $F(1,39) = 1.35$ , $p = 0.25$ , $\eta^2 = 0.034$ .

##### 2 Supplementary Figures

**Supplementary Figure 1.**
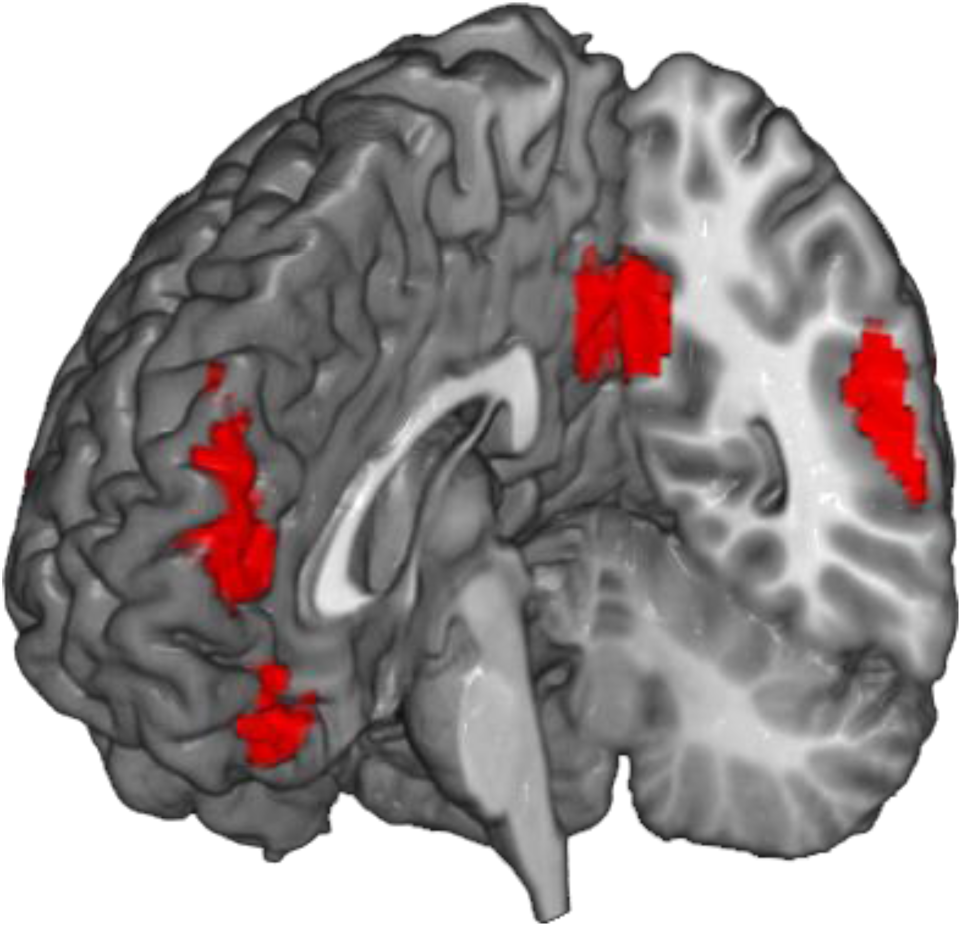
Selection of ROIs. To identify regions that are involved in mentalizing and theory of mind, but also show modulation of their activity by emotion, a conjunction of the neurosynth meta-analyses for the terms “theory of mind” (reverse inference map for regions *specifically* activated during theory of mind) and “emotion” (forward inference map for regions *consistently* activated by emotion) was computed. Overlap between the emotion and theory of mind networks was observed in left TPJ (k = 262, with only a few voxels found in right TPJ, k = 29) and DMPFC (k = 481), establishing these regions as prime candidates for investigations of the modulatory effects of emotion on mentalizing during trust decisions. Additional regions were identified in Posterior Cingulate Cortex (PCC, k = 314) and ventromedial PFC (k = 109), as well as bilateral superior temporal sulcus and inferior frontal gyrus (not shown).

###### 1.2 Emotion induction manipulation checks

**Supplementary Figure 2.**
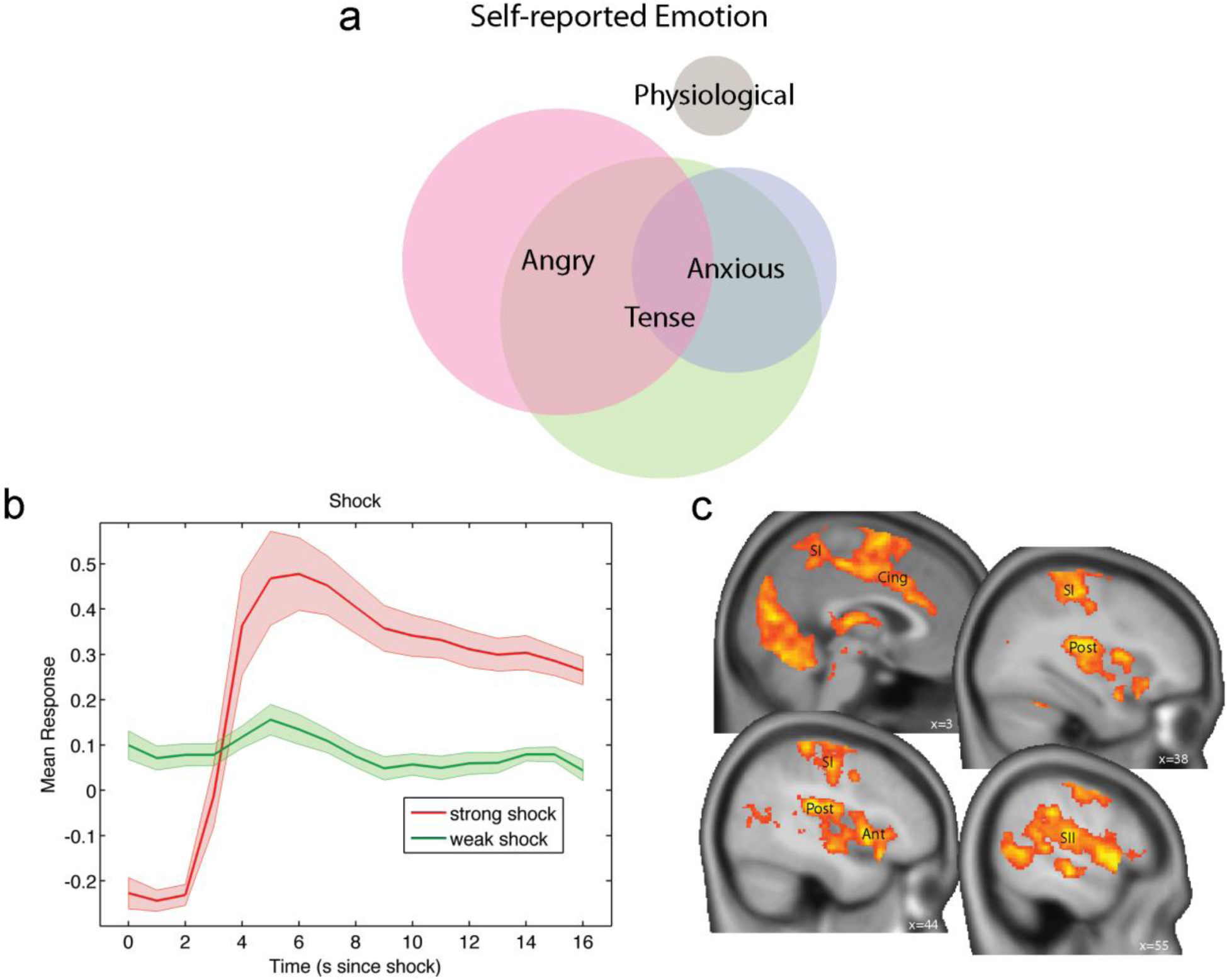
Manipulation checks. (a) Self-reported experience of emotion during threat blocks. Venn diagram illustrating proportions and overlap between self-reported emotional reactions to painful tactile stimulation. Note that subjects were able to report multiple emotions in response to the open-ended question about how they felt during threat blocks in an exit questionnaire. 95.12% of subjects indicated that they experienced aversive emotional arousal during pain blocks, such as tenseness and/or anger and/or anxiety. To illustrate the frequency of emotional reactions to painful tactile stimulation, subjects’ answers were binned into three emotion categories that best summarize their emotional responses: angry and annoyed responses were grouped into the “Angry” category (25), tense, stressed and surprised responses were grouped into the “Tense” category (27), and scared, nervous, helpless, and sad responses were grouped into the “Anxious” category (11). Most subjects reported angry (61%) and tense (66%) emotions while a minority also felt anxious (27%). A fourth category was termed “physiological” and best characterizes responses that reported physiological reactions related to aversive emotional arousal during painful blocks, such as “sweating” (7%). Of note, due to the open-ended nature of the questions, subjects were able to indicate more than one emotional reaction, as illustrated by the overlap between different sets in the Venn diagram. (b) The SCR response after receiving mildly painful electrical stimulation. SCR response after an *actually experienced* strong tactile stimulation (red line) shows a significant increase after about 4 seconds relative to the experience of a weak tactile stimulation, which leaves SCR almost unchanged (green line). (c) Administration of painful compared to just-noticeable tactile stimulation led to increased activity within key regions of the pain matrix, including primary (SI) and secondary (SII) somatosensory cortex, anterior (Ant) and posterior (Post) insula, as well as mid cingulate cortex (Cing). The contrast images reflect activation at the time of strong vs. weak tactile stimulation and are thresholded at p < 0.05 FWE-corrected at the cluster level (with a cluster-forming voxel-level threshold of p<0.001).

###### 1.3 Additional post-hoc inspection of the significant interactions between threat and game type within TPJ and amygdala

**Supplementary Figure 3.**
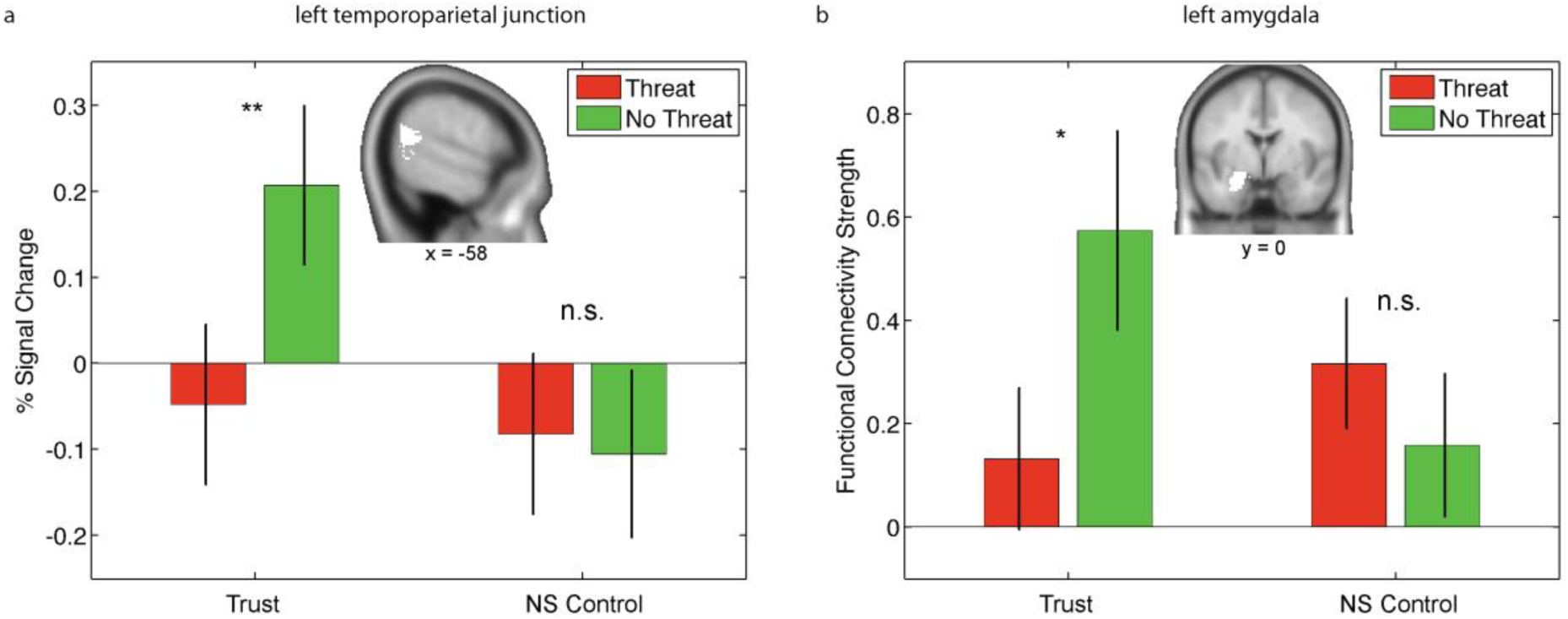
Additional post-hoc inspection of the significant interactions reported in the main paper within all voxels of independent TPJ and amygdala masks. To characterize the interaction patterns reported in the main paper, we extracted subject-specific regression coefficients (beta weights) from all voxels within the entire left TPJ (k = 1031) and amygdala (k = 439) masks and performed post-hoc statistical analyses in independent ROIs. (a) Post-hoc pairwise comparisons in the TPJ ROI revealed significantly greater choice-related activation during no-threat relative to threat when subjects made decisions in the trust task [t(40) = 2.769, p = 0.0085], but not when they made decisions in the control task [t(40) = -0.167, p = 0.8682]. (b) Post-hoc pairwise comparisons in the amygdala ROI revealed significantly greater choice-related functional connectivity during no-threat relative to threat when subjects made decisions in the trust task [t(40) = 2.097, p = 0.0424], but not when they made decisions in the control task [t(40) = -0.812, p = 0.4216]. These results confirm that the presence of threat during trust decisions led to a significant suppression of (a) TPJ activity and (b) TPJ-amygdala connectivity relative to the absence of threat. ** p < 0.01; * p < 0.05; n.s.: not significant

###### 1.4 Threat impacts domain general neural circuitry during both social and non-social decision making

**Supplementary Figure 4.**
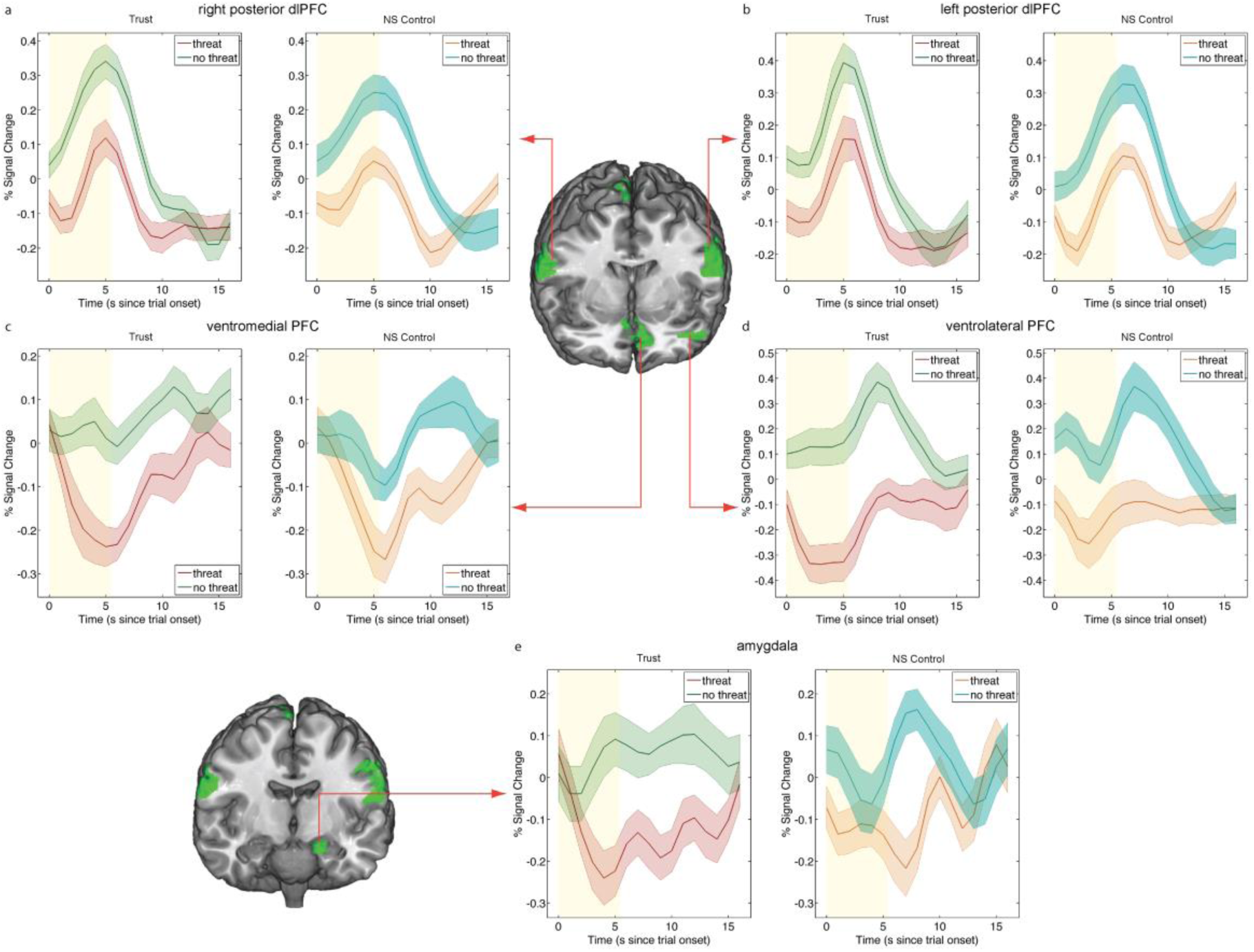
Main Effect of threat 1: Suppression of game type independent neural correlates of decision-making. We tested the main effect of threat on the neural correlates of decision-making independent of the game type (by including both trust and non-social decisions). This analysis revealed a domain-general network of regions showing significant threat-related reduction (no threat > threat) in choice-related activity during both trust and non-social control trials (see Supplementary Table 7a), consisting of bilateral posterior dlPFC [(a) right: 62, -6, 28, k = 1010 and (b) left: -62, -4, 18, k = 1901], left amygdala [(f) -24, -15, -23, k = 552], posterior paracentral lobule [not shown, 4 -36, 69, k = 887], and a large cluster in ventral anterior prefrontal cortex (k = 4082) that includes vmPFC [(c) -10, 44, -8] and left vlPFC [(d) -48, 41, -8]. Time courses illustrate suppression (a-e) due to threat during trust and non-social control decisions and are shown for both trust (left, threat shown in red, no threat shown in green) and control (right, threat shown in orange, no threat shown in aqua green) decisions in separate graphs. The 5.5-second choice period is displayed in yellow.

**Supplementary Figure 5.**
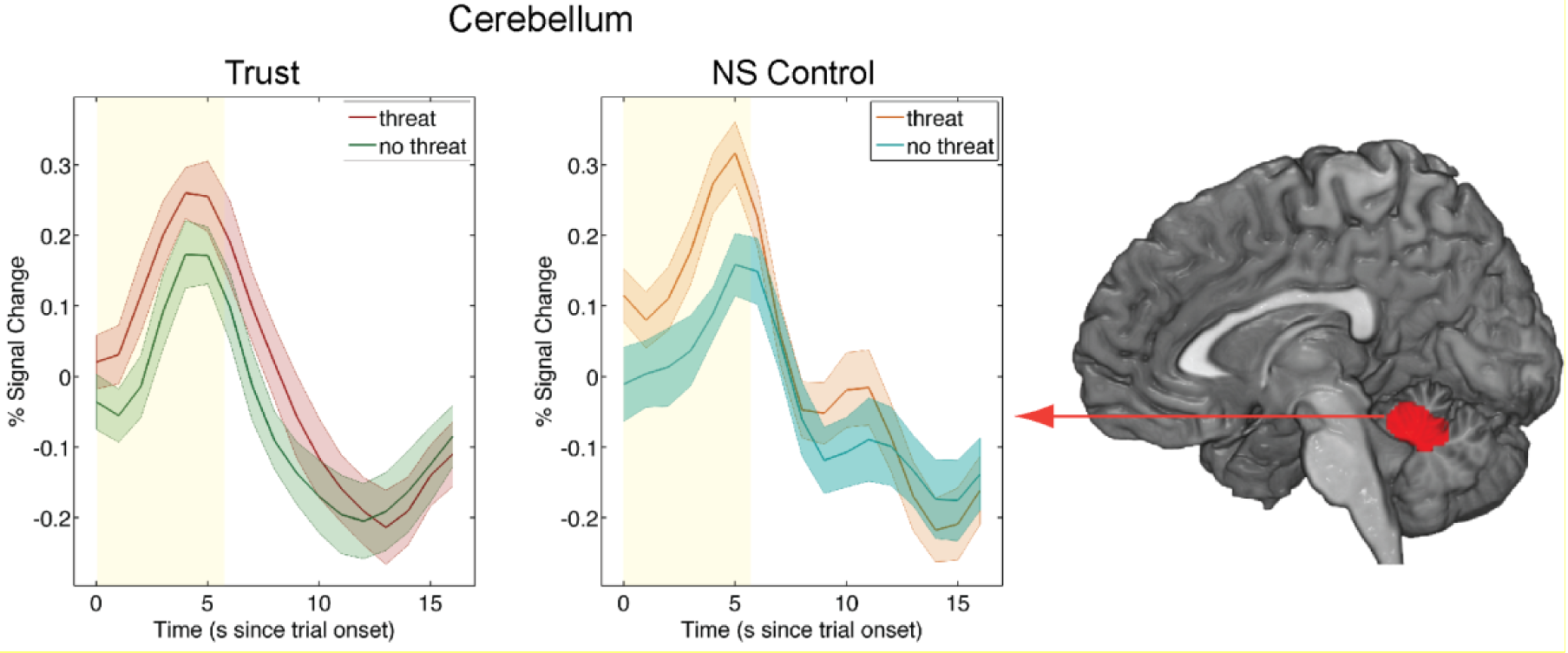
Main effect of threat 2: Enhancement of game type independent neural correlates of decision-making. We tested the main effect of threat on the neural correlates of decision-making independent of the game type (including both trust and non-social decisions). In addition to regions showing significant suppression of choice-related activity (Supplementary Figure 4, Supplementary Table 7a), we also found significant threat-related enhancement of activity (threat > no threat) during the decision phase (see Supplementary Table 7b) in the thalamus [(a) 18, -6, 1, k = 559] and cerebellum [(b) -4, -46, -24, k = 849]. Time courses illustrate enhancements (a-b) due to threat during trust and non-social control decisions and are shown for both trust (left, threat shown in red, no threat shown in green) and NS control (right, threat shown in orange, no threat shown in aqua green) decisions in separate graphs. The 5.5-second choice period is displayed in yellow.

#### 2. Supplementary Tables

**2.1 Supplementary Table 2.** Region of interest analyses investigating trust-specific neural correlates in regions associated with social cognition and valuation. (see Figure 2). (a) Regions that are preferentially involved in the trust game relative to the non-social control task when threat is absent. (b) Only the temporoparietal junction (TPJ) shows a significant interaction effect reflective of a change in the neural impact of threat during trust taking relative to the non-social control task. (c) Regions that show significant simple effects of threat during trust decisions.

| Structure | L/R | Cluster Size | Peak t | x | y | z | Peak p |
| --- | --- | --- | --- | --- | --- | --- | --- |
| *(a) Main effect: Trust (No Threat + Threat) > NS Control (No Threat + Threat)* |  |  |  |  |  |  |  |
| TPJ | left | 9 | 3.89 | -57 | -60 | 27 | 0.015 |
| ----- |  |  |  |  |  |  |  |
| *(b) Interaction: Trust (No Threat > Threat) – NS Control (No Threat > Threat)* |  |  |  |  |  |  |  |
| TPJ | left | 34 | 3.63 | -60 | -54 | 19 | 0.035 |
| Anterior Insula | left | 21 | 4.1 | -46 | -14 | -12 | 0.022 |
| ----- |  |  |  |  |  |  |  |
| *(c) Simple effect: Trust (No Threat > Threat)* |  |  |  |  |  |  |  |
| TPJ | left | 35 | 3.97 | -58 | -55 | 19 | 0.012 |
| vmPFC | bil. | 227 | 4.66 | 9 | 33 | -12 | 0.002 |
| Ventral striatum | left | 19 | 4.09 | -3 | 3 | -6 | 0.010 |

Activation clusters survive small volume correction for multiple comparisons based on FWE correction at the peak level for a priori regions of interest using masks created with reverse inference maps from relevant search terms on the neurosynth.org database. The left TPJ masks contained 1031 voxels; the dorsomedial and ventromedial PFC masks contained 3175 and 1327 voxels, respectively, the left ventral striatum mask contained 3239 voxel, and the anterior insula mask contained 13844 voxels.

**2.3 Supplementary Table 3.** Region of interest analyses investigating TPJ-amygdala connectivity patterns. (see Figures 3c-d). (a) TPJ-amygdala connectivity during decision-making in the trust relative to the control task *when threat is absent* and (b-c) the differential impact of threat on TPJ-amygdala connectivity in the trust task relative to the control task.

| Structure | L/R | Cluster Size | Peak t | x | y | z | Peak p |
| --- | --- | --- | --- | --- | --- | --- | --- |
| *(a) Main effect: No Threat (Trust + NS Control) > Threat (Trust + NS Control)* |  |  |  |  |  |  |  |
| amygdala | left | 14 | 3.78 | -28 | -6 | -14 | 0.008 |
| ----- |  |  |  |  |  |  |  |
| *(b) Interaction: Trust (No Threat > Threat) – NS Control (No Threat > Threat)* |  |  |  |  |  |  |  |
| amygdala | left | 12 | 3.68 | -26 | 0 | -23 | 0.012 |
| ----- |  |  |  |  |  |  |  |
| *(c) Simple effect: Trust (No Threat > Threat)* |  |  |  |  |  |  |  |
| amygdala | left | 23 | 4.03 | -28 | -6 | -14 | 0.004 |
| amygdala | left | 3 | 3.39 | -26 | 0 | -23 | 0.027 |
| ----- |  |  |  |  |  |  |  |
| *(d) Simple effect: Threat (NS Control > Trust)* |  |  |  |  |  |  |  |
| amygdala | left | 14 | 3.24 | -26 | 0 | -23 | 0.035 |

Activation clusters survive small volume correction for multiple comparisons based on FWE correction at the peak level for a priori regions of interest using anatomical masks created with reverse inference maps from relevant search terms on the neurosynth.org database. The left amygdala mask contained 439 voxels.

**2.4 Supplementary Table 4.** Region of interest analyses investigating brain-behavior relationships. ROI analyses investigating how the correlation between TPJ functional connectivity with its target regions and mean transfer rate is influenced by (a) game type in the absence of threat; (b) threat in the trust but not the NS control game; and (c) threat during trust decisions.

| Structure | L/R | Cluster Size | Peak t | x | y | z | Peak p |
| --- | --- | --- | --- | --- | --- | --- | --- |
| *(a) Main effect: Trust (No Threat + Threat) > NS Control (No Threat + Threat)* |  |  |  |  |  |  |  |
| STS (TPJ) | right | 30 | 3.75 | 64 | -43 | 4 | 0.034 |
| STS (TPJ) | left | 10 | 4.59 | -62 | -52 | -5 | 0.001 |
| DMPFC | bil. | 30 | 4.04 | -12 | 54 | 40 | 0.021 |
| DMPFC | bil | 9 | 3.91 | -3 | 38 | 48 | 0.037 |
| Amygdala | right | 16 | 3.49 | 28 | 2 | -20 | 0.024 |
| Anterior Insula | left | 245 | 5.16 | -51 | 21 | -6 | <0.001 |
| Anterior Insula | right | 525 | 4.5 | 56 | 18 | 1 | 0.006 |
| ----- |  |  |  |  |  |  |  |
| *(b) Interaction: Trust (No Threat > Threat) – NS Control (No Threat > Threat)* |  |  |  |  |  |  |  |
| STS (TPJ) | right | 7 | 4.73 | 64 | -43 | 6 | 0.001 |
| ----- |  |  |  |  |  |  |  |
| *(c) Simple effect: Trust (No Threat > Threat)* |  |  |  |  |  |  |  |
| STS (TPJ) | right | 31 | 5.63 | 64 | -43 | 6 | <0.001 |
| Anterior Insula | Left | 86 | 3.95 | -46 | 23 | -8 | 0.035 |

Activation clusters survive small volume correction for multiple comparisons based on FWE correction at the peak level for a priori regions of interest using masks created with reverse inference maps from relevant search terms on the neurosynth.org database. The right amygdala mask contained 492 voxels, the right TPJ mask contained 1416 voxels, the dmpFC mask contained 3175 voxels, the left anterior insula mask contained 13844 voxels, the right anterior insula mask contained 13871 voxels.

**2.5 Supplementary Table 5.**
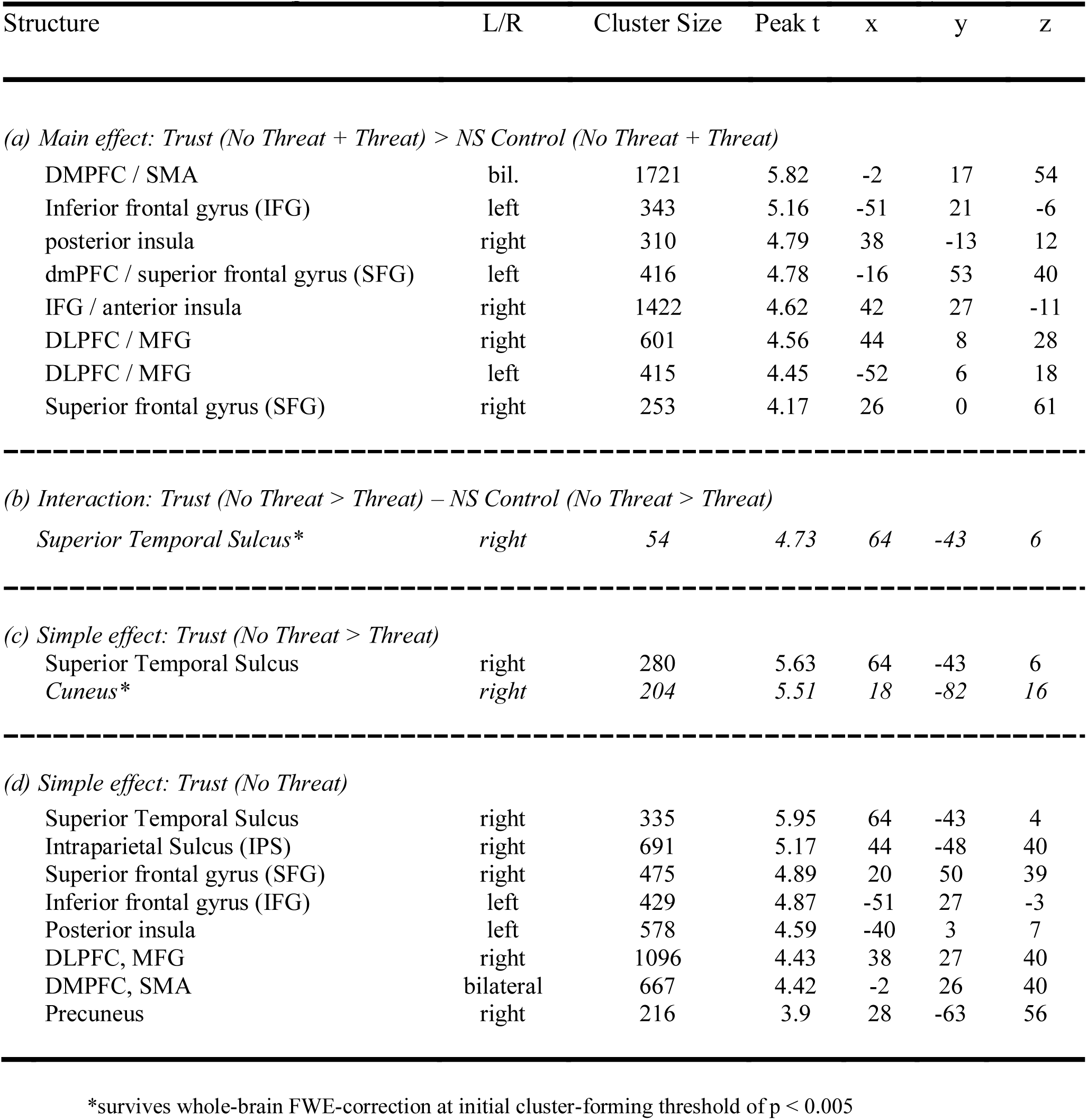
Whole-brain analyses investigating brain-behavior relationships. (see Figure 3). Results from whole-brain analysis investigating how the correlation between TPJ functional connectivity with its target regions and mean transfer rate is influenced by (a) game type in the absence of threat; (b) threat in the trust but not the NS control game; and (c) threat during trust decisions (p < 0.001, cluster size > 226, FWE corrected at cluster-level).

**2.6 Supplementary Table 6.**
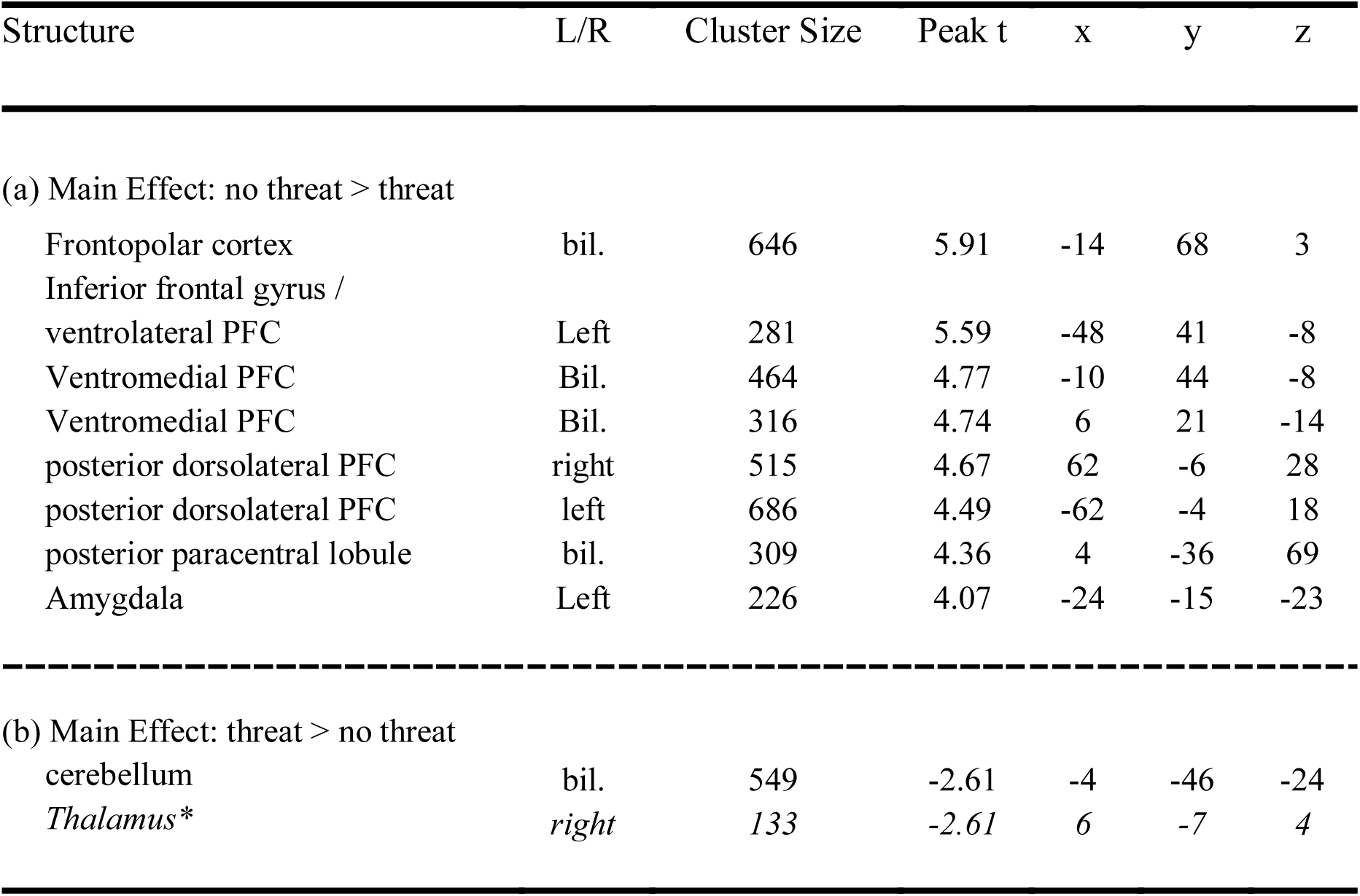
Whole-brain analyses investigating the main effect of threat. (see Supplementary Figure 4). Results from whole-brain analysis investigating the main effect of threat on neural activity during decision-making independent of game type (p < 0.001, cluster size > 226, FWE corrected at cluster-level).

## References

1. Delgado, M. R., Frank, R. H. & Phelps, E. A. Perceptions of moral character modulate the neural systems of reward during the trust game. Nat Neurosci 8, 1611–1618 (2005).

2. Bohnet, I. & Zeckhauser, R. Trust, risk and betrayal. Journal of Economic Behavior & Organization 55, 467–484 (2004).

3. Aimone, J. A., Houser, D. & Weber, B. Neural signatures of betrayal aversion: an fMRI study of trust. Proceedings of the Royal Society B: Biological Sciences 281, 20132127–20132127 (2014).

4. Fehr, E. On the Economics and Biology of Trust. Journal of the European Economic Association 7, 235–266 (2009).

5. Van Overwalle, F. Social cognition and the brain: a meta-analysis. Hum. Brain Mapp. 30, 829–858 (2009).

6. FeldmanHall, O., Raio, C. M., Kubota, J. T., Seiler, M. G. & Phelps, E. A. The Effects of Social Context and Acute Stress on Decision Making Under Uncertainty. Psychological Science 26, 1918–1926 (2015).

7. Mislin, A., Williams, L. V. & Shaughnessy, B. A. Motivating trust: Can mood and incentives increase interpersonal trust? Journal of Behavioral and Experimental Economics 58, 11–19 (2015).

8. Porcelli, A. J. & Delgado, M. R. Acute stress modulates risk taking in financial decision making. Psychological Science 20, 278–283 (2009).

9. Lerner, J. S., Small, D. A. & Loewenstein, G. Heart strings and purse strings: Carryover effects of emotions on economic decisions. Psychological Science 15, 337–341 (2004).

10. Cohn, A., Engelmann, J., Fehr, E. & Maréchal, M. A. Evidence for Countercyclical Risk Aversion: An Experiment with Financial Professionals. American Economic Review 105, 860–885 (2015).

11. Harlé, K. M. & Sanfey, A. G. Incidental sadness biases social economic decisions in the Ultimatum Game. Emotion 7, 876–881 (2007).

12. Loewenstein, G. Emotions in Economic Theory and Economic Behavior. 90, 426–432 (2000).

13. Phelps, E. A., Lempert, K. M. & Sokol-Hessner, P. Emotion and decision making: multiple modulatory neural circuits. Annu. Rev. Neurosci. 37, 263–287 (2014).

14. Lerner, J. S., Li, Y., Valdesolo, P. & Kassam, K. S. Emotion and decision making. Annu. Rev. Psychol. 66, 799–823 (2015).

15. Lindquist, K. A., Wager, T. D., Kober, H., Bliss-Moreau, E. & Barrett, L. F. The brain basis of emotion: A meta-analytic review. Behav Brain Sci 35, 121–143 (2012).

16. Pessoa, L. & Adolphs, R. Emotion processing and the amygdala: from a ‘low road’ to ‘many roads’ of evaluating biological significance. Nat Rev Neurosci 11, 773–783 (2010).

17. Kable, J. W. & Glimcher, P. W. The Neurobiology of Decision: Consensus and Controversy. Neuron 63, 733–745 (2009).

18. Rangel, A. & Hare, T. Neural computations associated with goal-directed choice. Current Opinion in Neurobiology 20, 262–270 (2010).

19. Kuhnen, C. M. & Knutson, B. The Influence of Affect on Beliefs, Preferences, and Financial Decisions. 46, 605–626 (2011).

20. Schulreich, S. et al. Music-evoked incidental happiness modulates probability weighting during risky lottery choices. Front. Psychol. 4, 981 (2014).

21. Phelps, E. A. in Neuroeconomics (eds. Glimcher, P. W., Fehr, E., Camerer, C. F. & Poldrack, R. A.) 233–250 (Elsevier, 2008).

22. Berg, J., Dickhaut, J. & McCabe, K. Trust, Reciprocity, and Social History. Games and Economic Behavior 10, 122–142 (1995).

23. McCabe, K., Houser, D., Ryan, L., Smith, V. & Trouard, T. A functional imaging study of cooperation in two-person reciprocal exchange. Proceedings of the National Academy of Sciences 98, 11832–11835 (2001).

24. Krueger, F. et al. Neural correlates of trust. Proceedings of the National Academy of Sciences 104, 20084–20089 (2007).

25. Baumgartner, T., Heinrichs, M., Vonlanthen, A., Fischbacher, U. & Fehr, E. Oxytocin shapes the neural circuitry of trust and trust adaptation in humans. Neuron 58, 639–650 (2008).

26. van den Bos, W., van Dijk, E., Westenberg, M., Rombouts, S. A. R. B. & Crone, E. A. What motivates repayment? Neural correlates of reciprocity in the Trust Game. Social Cognitive and Affective Neuroscience 4, 294–304 (2009).

27. Fett, A.-K. J., Gromann, P. M., Giampietro, V., Shergill, S. S. & Krabbendam, L. Default distrust? An fMRI investigation of the neural development of trust and cooperation. Social Cognitive and Affective Neuroscience 9, 395–402 (2014).

28. Bohnet, I., Greig, F., Herrmann, B. & Zeckhauser, R. Betrayal Aversion: Evidence from Brazil, China, Oman, Switzerland, Turkey, and the United States. American Economic Review 98, 294–310 (2008).

29. Kosfeld, M., Heinrichs, M., Zak, P. J., Fischbacher, U. & Fehr, E. Oxytocin increases trust in humans. Nature 435, 673–676 (2005).

30. Fehr, E. & Camerer, C. F. Social neuroeconomics: the neural circuitry of social preferences. Trends in Cognitive Sciences 11, 419–427 (2007).

31. Gromann, P. M. et al. Trust versus paranoia: abnormal response to social reward in psychotic illness. Brain 136, 1968–1975 (2013).

32. Sripada, C. S. et al. Functional neuroimaging of mentalizing during the trust game in social anxiety disorder. NeuroReport 20, 984–989 (2009).

33. Stanley, D. A. et al. Race and reputation: perceived racial group trustworthiness influences the neural correlates of trust decisions. Philosophical Transactions of the Royal Society B: Biological Sciences 367, 744–753 (2012).

34. Krueger, F., Grafman, J. & McCabe, K. Neural correlates of economic game playing. Philosophical Transactions of the Royal Society B: Biological Sciences 363, 3859–3874 (2008).

35. Saxe, R. & Kanwisher, N. People thinking about thinking peopleThe role of the temporo-parietal junction in ‘theory of mind’. NeuroImage 19, 1835–1842 (2003).

36. Behrens, T. E. J., Hunt, L. T. & Rushworth, M. F. S. The Computation of Social Behavior. Science 324, 1160–1164 (2009).

37. Ruff, C. C. & Fehr, E. The neurobiology of rewards and values in social decision making. Nature Reviews Neuroscience 15, 549–562 (2014).

38. Bogdan, R. & Pizzagalli, D. A. Acute stress reduces reward responsiveness: implications for depression. BPS 60, 1147–1154 (2006).

39. Grillon, C., Ameli, R., Foot, M. & Davis, M. Fear-potentiated startle: Relationship to the level of state/trait anxiety in healthy subjects. Biological Psychiatry 33, 566–574 (1993).

40. Schmitz, A. & Grillon, C. Assessing fear and anxiety in humans using the threat of predictable and unpredictable aversive events (the NPU-threat test). Nat Protoc 7, 527–532 (2012).

41. Westermann, R., Spies, K., Stahl, G. & Hesse, F. W. Relative effectiveness and validity of mood induction procedures: a meta analysis. Eur. J. Soc. Psychol. 26, 557–580 (1996).

42. Martin, M. On the induction of mood. Clinical Psychology Review 10, 669–697 (1990).

43. Starcke, K., Wolf, O. T., Markowitsch, H. J. & Brand, M. Anticipatory stress influences decision making under explicit risk conditions. Behavioral Neuroscience 122, 1352–1360 (2008).

44. Galvan, A. & Rahdar, A. The neurobiological effects of stress on adolescent decision making. Neuroscience 249, 223–231 (2013).

45. Shackman, A. J. et al. Anxiety selectively disrupts visuospatial working memory. Emotion 6, 40–61 (2006).

46. Robinson, O. J., Vytal, K., Cornwell, B. R. & Grillon, C. The impact of anxiety upon cognition: perspectives from human threat of shock studies. Front. Hum. Neurosci. 7, 203 (2013).

47. Young, L., Camprodon, J. A., Hauser, M., Pascual-Leone, A. & Saxe, R. Disruption of the right temporoparietal junction with transcranial magnetic stimulation reduces the role of beliefs in moral judgments. Proceedings of the National Academy of Sciences 107, 6753–6758 (2010).

48. Mitchell, J. P., Macrae, C. N. & Banaji, M. R. Dissociable medial prefrontal contributions to judgments of similar and dissimilar others. Neuron 50, 655–663 (2006).

49. Wicker, B. et al. Both of us disgusted in My insula: the common neural basis of seeing and feeling disgust. Neuron 40, 655–664 (2003).

50. Paulus, M. P. & Stein, M. B. An insular view of anxiety. BPS 60, 383–387 (2006).

51. Bellucci, G., Chernyak, S. V., Goodyear, K., Eickhoff, S. B. & Krueger, F. Neural signatures of trust in reciprocity: A coordinate-based meta-analysis. Hum. Brain Mapp. 1–17 (2016). doi:10.1002/hbm.23451

52. LeDoux, J. The amygdala. Current Biology 17, R868–74 (2007).

53. Ochsner, K. N., Silvers, J. A. & Buhle, J. T. Functional imaging studies of emotion regulation: a synthetic review and evolving model of the cognitive control of emotion. Annals of the New York Academy of Sciences 1251, E1–E24 (2012).

54. Engell, A. D., Haxby, J. V. & Todorov, A. Implicit Trustworthiness Decisions: Automatic Coding of Face Properties in the Human Amygdala. Journal of Cognitive Neuroscience 19, 1508–1519 (2007).

55. Winston, J. S., Strange, B. A., O’Doherty, J. & Dolan, R. J. Automatic and intentional brain responses during evaluation of trustworthiness of faces. Nat Neurosci 5, 277–283 (2002).

56. Weaver, B. & Wuensch, K. L. SPSS and SAS programs for comparing Pearson correlations and OLS regression coefficients. Behavior Research Methods 45, 880–895 (2013).

57. Yarkoni, T., Poldrack, R. A., Nichols, T. E., Van Essen, D. C. & Wager, T. D. Large-scale automated synthesis of human functional neuroimaging data. Nature Methods 8, 665–670 (2011).

58. Bartra, O., McGuire, J. T. & Kable, J. W. The valuation system: a coordinate-based meta-analysis of BOLD fMRI experiments examining neural correlates of subjective value. NeuroImage 76, 412–427 (2013).

59. Pessoa, L. Embedding reward signals into perception and cognition. 4, 1–8 (2010).

60. McLaren, D. G., Ries, M. L., Xu, G. & Johnson, S. C. A generalized form of context-dependent psychophysiological interactions (gPPI): A comparison to standard approaches. NeuroImage 61, 1277–1286 (2012).

61. Bohnet, I. & Zeckhauser, R. Trust, risk and betrayal. Journal of Economic Behavior & Organization 55, 467–484 (2004).

62. McCabe, K. A., Rigdon, M. L. & Smith, V. L. Positive reciprocity and intentions in trust games. Journal of Economic Behavior & Organization 52, 267–275 (2003).

63. Behrens, T. E. J., Hunt, L. T., Woolrich, M. W. & Rushworth, M. F. S. Associative learning of social value. Nature 456, 245–249 (2008).

64. Carter, R. M., Bowling, D. L., Reeck, C. & Huettel, S. A. A Distinct Role of the Temporal-Parietal Junction in Predicting Socially Guided Decisions. Science 337, 109–111 (2012).

65. Morishima, Y. et al. Linking Brain Structure and Activationin Temporoparietal Junctionto Explain the Neurobiology of Human Altruism. Neuron 75, 73–79 (2012).

66. Pessoa, L. Emotion and cognition and the amygdala: From “what is it?” to “what’s to be done?”. Neuropsychologia 48, 3416–3429 (2010).

67. Sander, D., Grafman, J. & Zalla, T. The Human Amygdala: An Evolved System for Relevance Detection : Reviews in the Neurosciences. Reviews in the Neurosciences 303–316 (2003). doi:10.1515/REVNEURO.2003.14.4.303","pf:authenticationStatus":"logged-in","pf:selectedLanguage":"English","pf:contentCategory":"nlm-article"}

68. Gromann, P. M. et al. Trust versus paranoia: abnormal response to social reward in psychotic illness. Brain 136, 1968–1975 (2013).

69. Stanley, D. A. et al. Race and reputation: perceived racial group trustworthiness influences the neural correlates of trust decisions. Philosophical Transactions of the Royal Society B: Biological Sciences 367, 744–753 (2012).

70. Hein, G., Morishima, Y., Leiberg, S., Sul, S. & Fehr, E. The brain’s functional network architecture reveals human motives. Science 351, 1074–1078 (2016).

71. McClure, E. B. et al. Abnormal attention modulation of fear circuit function in pediatric generalized anxiety disorder. Archives of General Psychiatry 64, 97–106 (2007).

72. Fusar-Poli, P. et al. Distinct Effects of Δ9-Tetrahydrocannabinol and Cannabidiol on Neural Activation During Emotional Processing. Arch. Gen. Psychiatry 66, 95–105 (2009).

73. Borgwardt, S. J. et al. Neural basis of Δ-9-tetrahydrocannabinol and cannabidiol: effects during response inhibition. Biological Psychiatry 64, 966–973 (2008).

74. O’Doherty, J. P., Buchanan, T. W., Seymour, B. & Dolan, R. J. Predictive Neural Coding of Reward Preference Involves Dissociable Responses in Human Ventral Midbrain and Ventral Striatum. Neuron 49, 157–166 (2006).

75. Wilkinson, D. & Halligan, P. The relevance of behavioural measures for functional-imaging studies of cognition. Nature Reviews Neuroscience 5, 67–73 (2004).

76. Pashler, H. Dual-task interference in simple tasks: data and theory. Psychological Bulletin (1994).

77. Bradley, M. M., Codispoti, M., Cuthbert, B. N. & Lang, P. J. Emotion and motivation I: Defensive and appetitive reactions in picture processing. Emotion 1, 276–298 (2001).

78. Yarkoni, T. & Braver, T. S. in The Handbook of Individual Differences in Cognition (eds. Gruszka, A., Matthews, G. & Szymura, B.) 87–108 (2010).

79. Brooks, A. M. et al. From Bad to Worse: Striatal Coding of the Relative Value of Painful Decisions. Front. Neurosci. 4, (2010).

80. Singer, T. et al. Empathic neural responses are modulated by the perceived fairness of others. Nature 439, 466–469 (2006).

81. Fareri, D. S., Chang, L. J. & Delgado, M. R. Computational Substrates of Social Value in Interpersonal Collaboration. Journal of Neuroscience 35, 8170–8180 (2015).

82. Pruessmann, K. P., Weiger, M., Scheidegger, M. B. & Boesiger, P. SENSE: Sensitivity encoding for fast MRI. Magnetic Resonance in Medicine 42, 952–962 (1999).

83. Henson, R. N. A., Price, C. J., Rugg, M. D., Turner, R. & Friston, K. J. Detecting latency differences in event-related BOLD responses: application to words versus nonwords and initial versus repeated face presentations. NeuroImage 15, 83–97 (2002).

84. Rilling, J. K. & Sanfey, A. G. The Neuroscience of Social Decision-Making. Annu. Rev. Psychol. 62, 23–48 (2011).

85. Carter, R. M. & Huettel, S. A. A nexus model of the temporal– parietal junction. Trends in Cognitive Sciences 17, 328–336 (2013).

86. Levy, D. J. & Glimcher, P. W. The root of all value: a neural common currency for choice. Current Opinion in Neurobiology 1–12 (2012). doi:10.1016/j.conb.2012.06.001

87. Gläscher, J. Visualization of Group Inference Data in Functional Neuroimaging. Neuroinform 7, 73–82 (2009).

88. Gitelman, D. R., Penny, W. D., Ashburner, J. & Friston, K. J. Modeling regional and psychophysiologic interactions in fMRI: the importance of hemodynamic deconvolution. NeuroImage 19, 200–207 (2003).

